# Prognostic importance of splicing-triggered aberrations of protein complex interfaces in cancer

**DOI:** 10.1101/2024.05.06.592695

**Authors:** Khalique Newaz, Christoph Schaefers, Katja Weisel, Jan Baumbach, Dmitrij Frishman

**Affiliations:** Institute for Computational Systems Biology and Center for Data and Computing in Natural Sciences, Universität Hamburg, 22761, Hamburg, Germany; Department of Oncology, Hematology and Bone Marrow Transplantation with Division of Pneumology, Universitätsklinikum Hamburg-Eppendorf, 20251, Hamburg, Germany; Department of Mathematics and Computer Science, University of Southern Denmark, Odense, Denmark; Department of Bioinformatics, School of Life Sciences, Technical University of Munich, 85354, Freising, Germany

**Keywords:** alternative splicing, protein complexes, exon-exon interactions, cancer prognosis

## Abstract

Aberrant alternative splicing (AS) is a prominent hallmark of cancer. AS can perturb protein-protein interactions (PPIs) by adding or removing interface regions encoded by individual exons. Identifying prognostic exon-exon interactions (EEIs) from PPI interfaces can help discover AS-affected cancer-driving PPIs that can serve as potential drug targets. Here, we assessed the prognostic significance of EEIs across 15 cancer types by integrating RNA-seq data with three-dimensional (3D) structures of protein complexes. By analyzing the resulting EEI network we identified patient-specific perturbed EEIs (i.e., EEIs present in healthy samples but absent from the paired cancer samples or vice versa) that were significantly associated with survival. We provide the first evidence that EEIs can be used as prognostic biomarkers for cancer patient survival. Our findings provide mechanistic insights into AS-affected PPI interfaces. Given the ongoing expansion of available RNA-seq data and the number of 3D structurally-resolved (or confidently predicted) protein complexes, our computational framework will help accelerate the discovery of clinically important cancer-promoting AS events.

## Introduction

Failure to identify patients with aggressive cancer and then to intervene in a timely manner dramatically lowers life expectancy of affected individuals (1). Molecular survival analysis aims to identify prognostic biomarkers, i.e., molecules whose abundance correlates with the survival rates of patients. Identification of such molecular prognostic biomarkers is crucial for stratification of patient groups according to their survival risk as well as for discovering potential anti-cancer drug targets.

Numerous studies have used molecular and clinical data from public databases, such as The Cancer Genome Atlas (TCGA) (2), to perform molecular survival analysis (3; 4; 5; 6; 7). One such recent study evaluated genome-wide associations between patient survival and six different types of molecular profiles: gene expression, protein expression, point mutations, copy number alterations, DNA methylation, and microRNA expression, across 33 cancer types (8). The study found that the number of prognostic molecules varied across cancer types, highlighting that different cancer types have different molecular prognostic biomarkers due to the difference in the underlying mechanisms of pathogenesis.

Since proteins often function by interacting with each other to form protein-protein interactions (PPIs), one promising direction of anti-cancer drug development has been to target PPIs (9). For example, GDC-0917 is a drug that targets the interaction between the Caspase-9 protein and X-linked inhibitor of apoptosis protein, and is in phase I trials for treatment of solid cancers (10). Aberrant PPIs have been related to cancer development (11; 12) and PPI-level differences have been reported among cancer types (13; 14), which could be due to the underlying differences in the patterns of cancer type-related mutations (15) or alternative splicing (AS) (16). For example, Climente-González et al. (17) identified a number of AS events that perturb PPIs across 11 cancer types. Furthermore, a recent study evaluated associations between patient survival and AS events across 31 cancer types (7). Although the prognostic value of individual AS events has been explored, a comprehensive large-scale investigation of AS-related PPI perturbations with cancer prognostic value has so far been lacking.

Here, we performed a pan-cancer analysis to evaluate the prognostic value of exon-exon interaction (EEI) interfaces of protein complexes across 15 cancer types that had enough paired (healthy and cancer) patient samples and transcriptomics data in TCGA. RNA-seq data were integrated with three-dimensional (3D) structures of heterodimeric protein complexes from the Protein Data Bank (PDB) (18). The number of perturbed EEIs (i.e., EEIs present in a healthy sample but absent from the paired cancer sample or vice versa) associated with patient survival exhibited strong variation across cancer types. Importantly, we show that many of such perturbed EEIs can be used as potential prognostic survival biomarkers (Figure 1). For example, two of the identified EEIs possessing prognostic value in breast cancer constitute an important part of the interaction interface in the go-ichi-ni-san (GINS) protein complex, which has been previously related to BRCA prognosis (19; 20). Our findings can not only unravel novel prognostic survival biomarkers, but can also be mined to gain insights into the interface regions of protein complexes that can potentially be targeted by anti-cancer drugs.

**Fig. 1:**
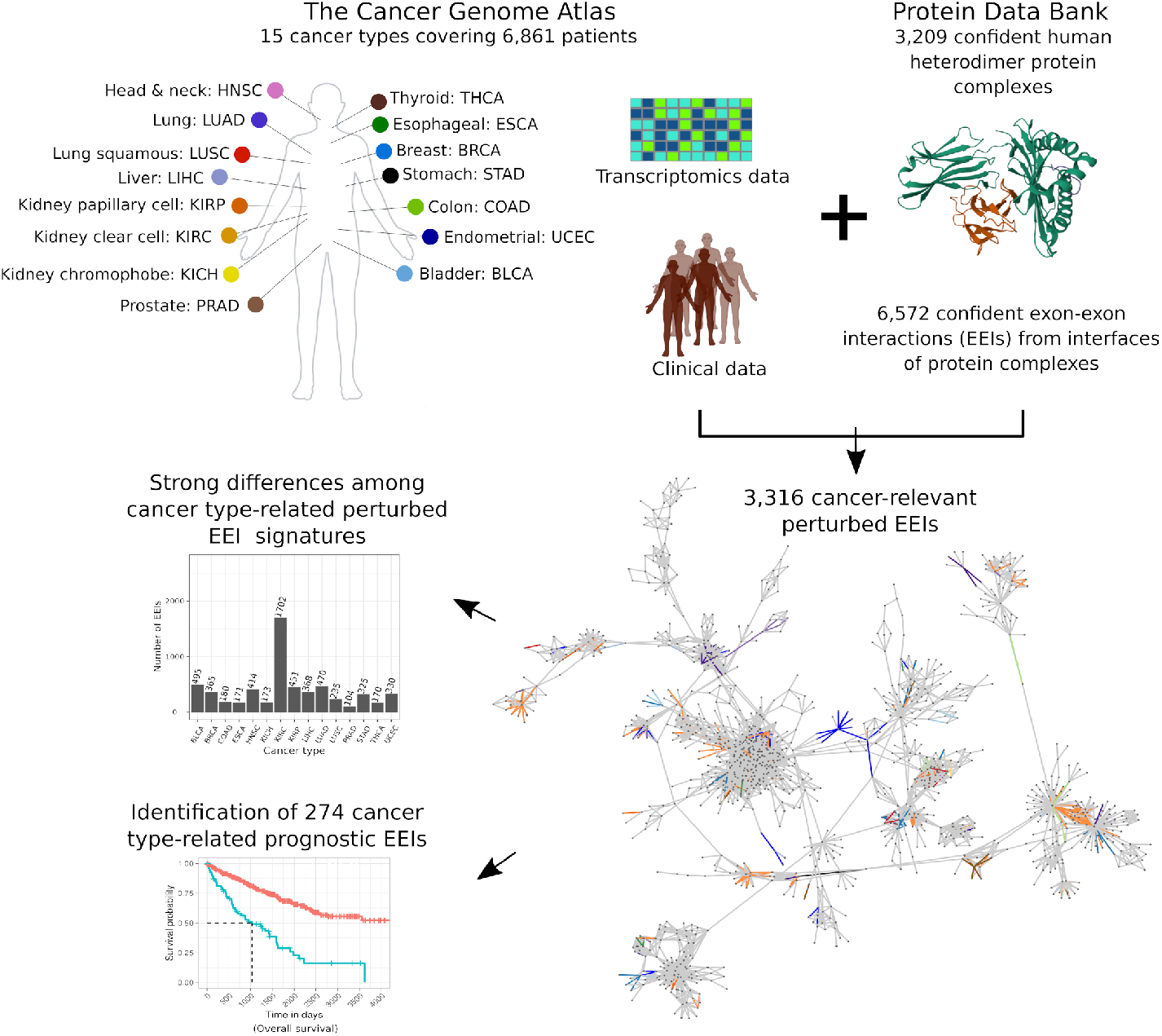
Overview of the study. Transcriptomics and clinical data from The Cancer Genome Atlas were integrated with structural protein complex data from the Protein Data Bank to identify cancer-relevant perturbed exon-exon interactions (EEIs), which were then used to derive cancer type-related EEI signatures and prognostic biomarkers.

## Data and methods

### Transcriptomics data

We identified primary solid tumor (cancer) and patient-matched control (healthy) samples in the TCGA database associated with those 15 cancer types that had at least 10 patients each (Table 1). To perform survival analysis for each cancer type (Section 2.6), we considered patients with at least one cancer sample and overall survival information. For the total of 7,547 TCGA samples (across 6,861 patients) obtained using the above two procedures, we downloaded the corresponding RNA-seq reads (bam files) from TCGA that were aligned to the Genome Reference Consortium Human Build 38.

**Table 1.** Patient counts across 15 cancer types.

| Cancer type | TCGA cancer code | # of patients with paired samples | # of patients with at least one cancer sample and overall survival information |
| --- | --- | --- | --- |
| Bladder urothelial carcinoma | BLCA | 19 | 403 |
| Breast invasive carcinoma | BRCA | 113 | 1093 |
| Colon adenocarcinoma | COAD | 41 | 455 |
| Esophageal carcinoma | ESCA | 13 | 184 |
| Head and neck squamous cell carcinoma | HNSC | 43 | 519 |
| Kidney chromophobe | KICH | 25 | 65 |
| Kidney renal clear cell carcinoma | KIRC | 72 | 533 |
| Kidney renal papillary cell carcinoma | KIRP | 32 | 289 |
| Liver hepatocellular carcinoma | LIHC | 50 | 369 |
| Lung adenocarcinoma | LUAD | 58 | 507 |
| Lung squamous cell carcinoma | LUSC | 51 | 495 |
| Prostate adenocarcinoma | PRAD | 52 | 495 |
| Stomach adenocarcinoma | STAD | 33 | 406 |
| Thyroid carcinoma | THCA | 59 | 505 |
| Uterine corpus endometrial carcinoma | UCEC | 23 | 543 |

Given a bam file, raw exon expression was quantified as follows. First, we downloaded the exon annotation from the Ensembl database (21). Second, we employed the testPairedEndBam function from Rsamtools R package to differentiate between the bam files containing paired-end and single reads. For the former, raw exon counts were quantified by featureCounts (22) using the command *featureCounts –p –O –t exon –g exon id –F GTF –T 8 –a EnsemblExonAnnotation*, while for the latter we used the command *featureCounts –O –t exon –g exon id –F GTF –T 8 –a EnsemblExonAnnotation*.

For each cancer type, given the raw expression values from each sample (cancer or healthy) of every patient, we removed technical bias in samples using the Trimmed Mean of M-values method (TMM) (23) to obtain the normalized exon counts, which were then converted into counts-per-million (CPM) values. An exon was defined to be expressed if the corresponding CPM value was greater than a chosen threshold. Six CPM value thresholds were tested in this work: 0.05, 0.1, 0.2, 0.5, 1, and 2.

### Dataset of heterodimeric protein complexes with known 3D structures and annotated exons

We identified 6,670 reviewed human proteins from the UniProt database (24) (downloaded in Jan. 2022) with the total of 41,330 known 3D structures available in the Crystallographic Information Files of the PDB, where each protein had at least one PDB structure determined either by X-ray crystallography or electron microscopy with better than 3Å resolution, or by nuclear magnetic resonance. To capture potential heterodimers, we selected PDB structures that contained at least two different co-crystallized proteins represented in distinct protein chains. If multiple alternative structures were available for a given pair of co-crystallized proteins, we selected the PDB entry with the highest number of structurally resolved amino acids. The above two selection criteria resulted in 13,967 heterodimers across 2,349 PDB entries covering 2,882 proteins.

To map exon information onto proteins, we matched UniProt IDs and Ensembl transcripts using the biomaRt R package. Out of many alternative transcripts of a given gene, we picked the one that matched exactly the corresponding UniProt protein based on pairwise global alignment. Heterodimers in which one or both proteins did not have such transcripts were excluded from further consideration. This procedure yielded 13,240 heterodimers across 2,088 PDB entries covering 2,660 proteins. For each of the 2,660 proteins, we mapped exon coordinates onto their amino acid sequences according to the Ensembl GTF file (“Homo sapiens.GRCh38.105.gtf.gz”).

Subsequently, we kept for further analyses only those proteins for which unambiguous correspondence between the amino acid sequences provided in the UniProt and the PDB could be established according to the Structure Integration with Function, Taxonomy, and Sequence (SIFTS) database (25). Our final dataset contained 13,190 heterodimers across 2,064 PDB IDs covering 2,649 proteins.

### Definitions of exon-exon interactions

For each pair of co-crystallized proteins, we evaluated whether any two exons from these two proteins interact, i.e., form an exon-exon interaction (EEI), using three alternative approaches: contact-based, energy-based, and evolution-based.

#### Contact-based approach

Two exons were considered to interact if there was at least one residue-residue contact between them. Contacts between any two residues were defined based on the spatial distance of 6Å between any of their heavy atoms, as is typically done to obtain biologically relevant protein contact maps (26; 27; 28; 29). The above procedure resulted in an EEI network (EEIN) with 16,012 EEIs among 9,442 exons.

#### Energy-based approach

We used PISA (Protein Interfaces, Surfaces, and Assemblies; https://www.ebi.ac.uk/msd-srv/prot_int/cgi-bin/piserver) (30) to evaluate whether an interface between two co-crystallized proteins is biologically relevant. PISA computes a solvation free energy gain upon interface formation between two proteins and then determines the probability of observing the same solvation free energy gain by chance when a random set of atoms (with the same area as that of the interface) is picked from the same two proteins surfaces not participating in the interface. PISA considers an interface to be biologically relevant if the corresponding probability is less than 0.5. For each PISA-defined biologically relevant interface, we considered each pair of interface exons originating from the two proteins as an EEI and thus obtained an EEIN with 18,619 EEIs among 6,946 exons.

#### Evolution-based approach

We used EPPIC (Evolutionary Protein-Protein Interface Classifier) (31) to evaluate whether an interface between two co-crystallized proteins is biologically relevant based on a combination of evolutionary information and surface area of the interface. EPPIC characterizes an interface between two proteins as biologically relevant if the interface surface area is greater than 2,200 Å^2^, while it characterizes an interface to be an artifact of crystallization if the interface surface area is smaller than 440 Å^2^. For any interface with an area between 400 Å^2^ and 2,200 Å^2^, EPPIC assesses the evolutionary conservation of the corresponding protein sequences to characterize whether the interface is biologically relevant or not. For each EPPIC-defined biologically relevant interface, we considered each pair of interface exons originating from the two proteins as an EEI and thus obtained an EEIN with 16,991 EEIs among 5,975 exons.

### Global and sample-specific EEINs at different confidence levels

We employed three global EEINs with a varying degree of confidence and coverage: (1) a high confidence network (NETHIGH) with the smallest coverage having 6,572 edges (*∼*25%) supported by all three EEI definition approaches (Section 2.3), (2) a medium confidence network (NETMEDIUM) with 18,418 edges (*∼*69%) supported by at least two EEI definition approaches, and (3) a low confidence network (NETLOW) with the largest coverage having 26,632 edges (100%) supported by at least one EEI definition approach (Figure 2).

**Fig. 2:**
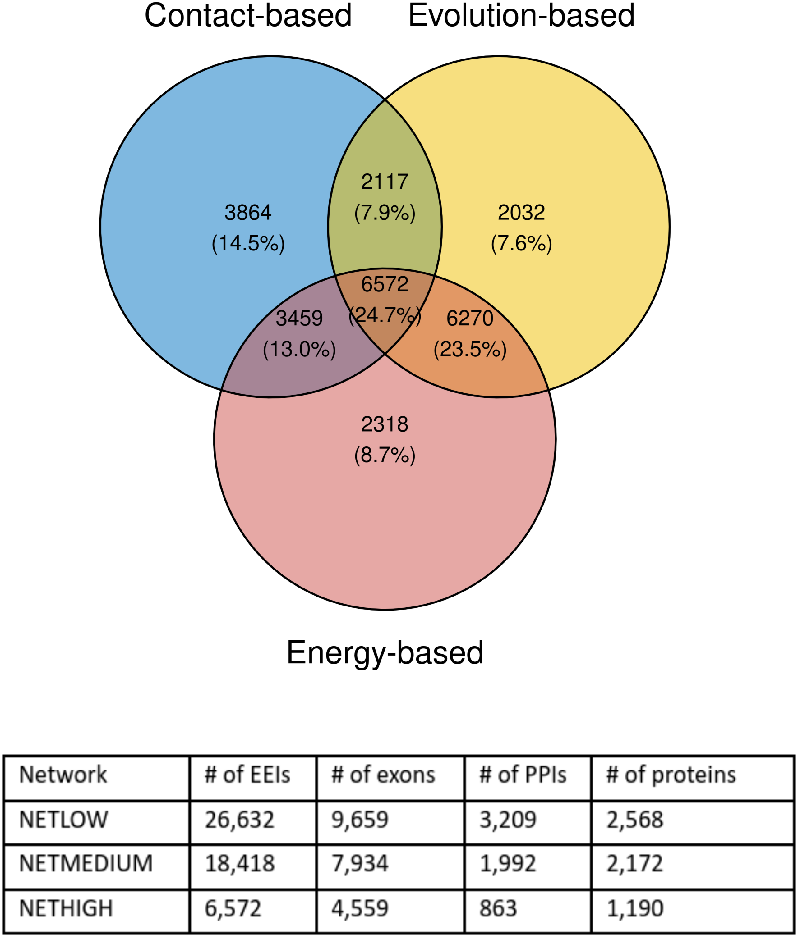
Details of the global EEINs at different confidence levels used in this study. The top panel outlines the overlap of the three global EEINs corresponding to the three EEI definition approaches. Given an area of the Venn diagram, the two numbers represent the numbers of edges and the percentage of edges over all edges (26,632) in the union of the three networks, respectively. The bottom panel shows the details of the network sizes of the final three global EEINs with different confidence and coverage.

Based on the exon expression data associated with a patient sample (cancer or healthy), sample-specific EEINs were derived as induced subgraphs of a global EEIN (i.e., any of NETHIGH, NETMEDIUM, or NETLOW) by considering all sample-specific expressed exons (Section 2.1) present in the EEIN along with the EEIs between them.

### Patient- and cancer type-related categories of exon-exon interactions

For each patient, we compared cancer and healthy sample-specific EEINs to identify three edge types in a global EEIN: (a) gained edges that were present in the cancer sample but were absent from the healthy sample, (b) lost edges that were present in the healthy sample but were absent from the cancer sample, and (c) non-perturbed edges that were either absent from both healthy and cancer samples or were present in both the cancer and healthy samples. These three types of edges are exemplified in Figure 3: (a) the edge {H,B} is a gained edge for patient 2 because it is absent from the healthy sample but is present in the paired cancer sample, (b) the edge {A,B} is a lost edge for patient 1 because it is present in the healthy sample but is absent from the paired cancer sample, (c) the edge {B,C} is a non-perturbed edge for patient 1 because it is present in both the healthy and cancer samples. Similarly, the edge {B,C} is a non-perturbed edge in patient 3 because it is absent from both the corresponding healthy and cancer samples.

**Fig. 3:**
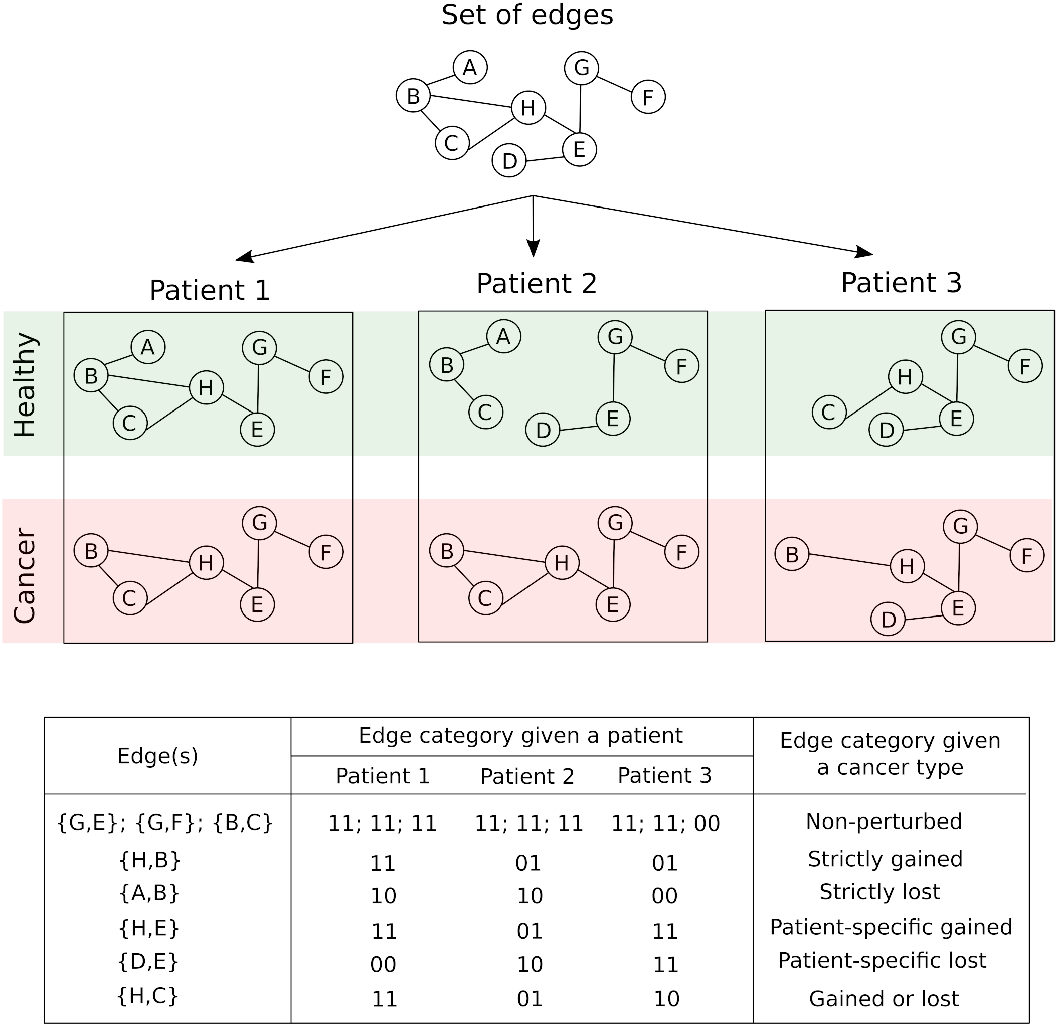
Partitioning of edges into edge categories for a given patient or cancer type, illustrated on a dummy set of eight edges. For each patient, three edge categories were defined: (a) gained edges, denoted “01”, which are not present in the healthy sample but are present in the paired cancer sample (b) lost edges, denoted “10”, which are present in the healthy sample but are absent from the paired cancer sample, and (c) non-perturbed edges, denoted either “00” or “11”, which are either absent from both healthy or cancer samples or present in both healthy or cancer samples. For each cancer type, we grouped the edges into six categories (last column in the table of this figure).

For each cancer type, given the gained, lost, and non-perturbed edges defined for every patient, we further defined the following six edge types: (a) non-perturbed edges that were not perturbed (“00” or “11” in Figure 3) in any of the considered patients, (b) strictly gained edges that were gained (“01” in Figure 3) in at least two patients but were not perturbed in any of the other patients, (c) strictly lost edges that were lost (“10” in Figure 3) in at least two patients but were not perturbed in any of the other patients, (d) patient-specific gained edges that were gained in exactly one patient but were not perturbed in any of the other patients, (e) patient-specific lost edges that were lost in exactly one patient but were not perturbed in any of the other patients, and (f) gained or lost edges that were lost in at least one patient and were gained in at least one other patient. An edge was categorized to be perturbed if it belonged to any of the five edge types (b) through to (f).

### Survival analysis

Given a cancer type with all of the corresponding patient samples, a particular edge connecting two exons, and an exon expression threshold, we investigated whether the presence (both exons expressed) or absence (one or both exons not expressed) of that edge is informative of patient survival rates. To ensure sufficient sample size, we only considered the edge if it partitioned patients into two groups with at least 10 patients in each group, such that the edge was present in all of the patients in one group while it was absent from all of the patients in the other group. We assessed whether the edge’s presence or absence is statistically significantly associated with the distinction between the patients with high vs. low survival rate by comparing the corresponding Kaplan-Meier survival curves and computing the logrank *p*-value. We considered an edge to be statistically significantly associated with the patient survival rate if the corresponding logrank *p*-value was smaller than or equal to 0.05.

### Genes significantly associated with patient survival

Associations between the mRNA expression of individual genes and patient survival were established using the data provided in the Supplementary Table S1 of the publication by Smith and Sheltzer (8), who correlated gene expression levels with clinical outcomes using the Cox proportional hazards regression. Only those 16,340 genes were considered that mapped to UniProt IDs as per the biomaRt package in R. As recommended by the authors, a gene was considered to be statistically significantly associated with patient survival of a given cancer type if its corresponding Z-score, calculated by dividing the regression coefficient by its standard error, was smaller than -1.96 or greater than 1.96, both of which correspond to a *p*-value of less than 0.05. Given a cancer type, we computed the overlap between the genes showing statistically significant associations with patient survival and any other given set of genes using the hypergeometric test. Here, we used all 16,340 genes as the background.

### Alternative splicing events significantly associated with patient survival

Associations between alternative splicing (AS) events (described in the SpliceSeq (32) and SpIAdder (33) resources) and patient survival were obtained from the study of Zhang et al. (34), which reported cancer type-related AS events that are significantly (*p*-value < 0.05) correlated with clinical outcomes according to the Cox proportional hazards regression. To obtain the number of cancer type-related AS events, the column named “SpliceSeq ASEs” was considered, while to obtain the number of AS events significantly associated with patient survival, the column named “SpliceSeq SASEs OS” was considered from the Supplementary Table S1 in Zhang et al. (34). The “SpliceSeq SASEs OS” column contained information for all cancer types considered in our study except PRAD and THCA.

### KEGG pathway enrichment

We used the KEGGREST R package to map KEGG pathways (35) to exons. To do this, we annotated each exon of a gene with all of the gene-related KEGG pathways. Note that we focus on the entire KEGG pathway and not on the individual KEGG modules which, if applicable, could be part of one or multiple KEGG pathways. Thus, our KEGG pathway enrichment analysis can only outline which pathways were enriched and can not pinpoint the enriched module(s) within a pathway. We focus only on those pathways that were associated with at least two exons in a given background EEIN. Given a set of exons, we measured the enrichment of each considered KEGG pathway individually using the hypergeometric test, which measures the probability (*p*-value) of getting the same or higher number of occurrences of a given KEGG pathway in another set of exons (with the same number of exons as the original set) that is randomly selected from a background set of exons. The background set of exons was constituted by all exons of the background EEIN that were annotated by at least one KEGG pathway. Correction for testing multiple pathways was performed by the Benjamini-Hochberg procedure, and the *p*-values were adjusted to obtain the corresponding *q*-values. We identified a pathway to be statistically significantly enriched if its *q*-value was less than or equal to 0.05.

### Cancer survival biomarker identification

A given edge was considered a potential survival biomarker if it showed stronger survival association (Section 2.6) on real data than on random data. Randomizations were performed using two alternative approaches. First, we kept the associations of survival times with patients the same as in the real data, but randomized the association of expressed edges to patients, while keeping the total number of patients in which a given edge is expressed the same as in the real data. Second, we kept the associations of expressed edges with patients the same as in the real data but randomized the associations of survival times with patients. Survival analysis using both randomization procedures was repeated 100 times, resulting in 100 random logrank *p*-values. An edge was considered to be a potential survival biomarker if (1) its actual logrank *p*-value was less than 0.05 and (2) all of the random *p*-values (for each of the two randomization tests) were greater than the actual logrank *p*-value.

## Results and discussion

### Identification of cancer-relevant perturbed edges

As outlined in Section 2.4, we created three global EEINs with varying confidence levels – NETHIGH, NETMEDIUM, and NETLOW. To define global EEINs, we used three EEI definition approaches: contact-based, energy-based, and evolution-based (Section 2.3). While the energy-based and evolution-based approaches potentially consider known characteristics of PPI interfaces to detect EEIs in co-crystallized protein complexes, the contact-based approach defines an interaction between two exons if there is at least one residue-residue contact between them. Thus, the contact-based approach is a relatively weaker EEI definition approach than the other two. For this reason, instead of just using each of three EEI definition approaches individually to define global EEINs, we focused on the EEIs confirmed by all three approaches to define global EEINs at three different confidence levels as listed above. Nevertheless, to validate the robustness of our key results, we repeated our study using a more stringent cutoff of at least five (instead of at least one) residue-residue contact for the contact-based EEI definition (Supplementary Section S1) and obtained qualitatively similar results. The contact-based EEI definition based on at least one residue-residue contact was used in this study.

Given a cancer type, for each perturbed edge (i.e., EEI) we performed survival analysis based on all corresponding cancer samples (Section 2.6). This procedure was applied to each of the 15 cancer types individually and all edge perturbations significantly (*p*-value ≤ 0.05) correlated with survival in at least one cancer type were considered to be cancer-relevant. Here, we used *p*-values from survival analysis as proxy scores to identify cancer-relevant perturbed EEIs. Hence, we did not perform a multiple testing correction as it would not change the relative scores of the EEIs. Nevertheless, to validate the robustness of our key results, we repeated survival analysis using an alternative *p*-value cutoff of 0.01 (Supplementary Section S2) and obtained qualitatively similar results. The *p*-value cutoff of 0.05 was used in this study.

Note that the identification of edges involved in cancer-relevant perturbations relies on the definition of exon expression (i.e., normalized CPM value, Section 2.1). To define whether a gene is expressed or not, CPM values ranging from 0.5 to 2 were used in previous studies (36; 37). We performed survival analysis using six different exon expression thresholds (Section 2.1) and found that the CPM value threshold of 0.5 resulted in the maximum number of cancer-relevant perturbed edges (CRPEs). As seen in Figure 4, the number of CRPEs first increases for the expression value thresholds in the range from 0.05 to 0.5, but then decreases with the further increase of expression value threshold from 0.5 to 2. These results correlate with the numbers of perturbed edges. Thus, although the numbers of edges in the paired control and cancer samples decrease along the CPM values of 0.05, 0.1, 0.2, 0.5, 1, and 2 (as expected), the number of perturbed edges first increases for the expression values from 0.05 to 0.5 and then decreases for the expression values from 0.5 to 2 (Figure 4). See Supplementary Section S3 for a detailed discussion of the dependence between the number of edges and the expression value threshold.

**Fig. 4:**
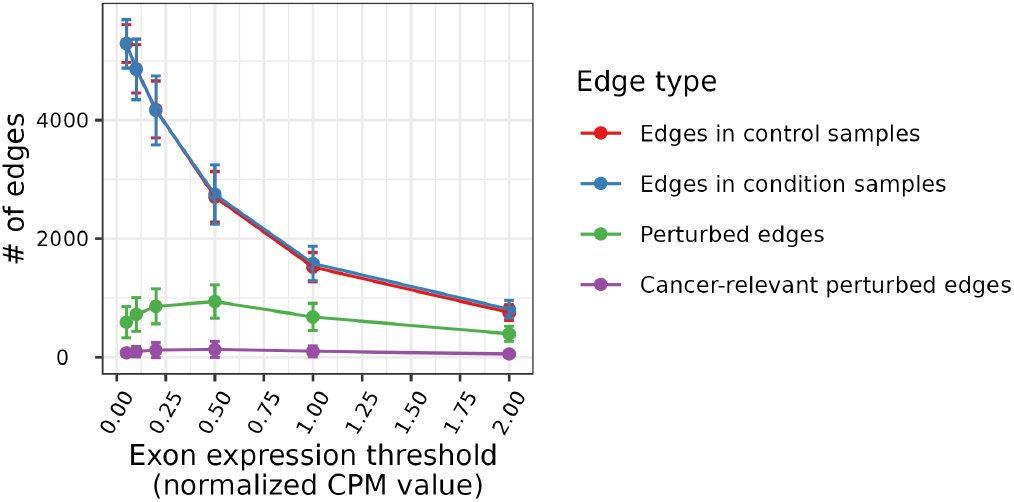
Dependence of the number of edges on the choice of expression value threshold (results shown for NETHIGH; results for NETMEDIUM and NETLOW were qualitatively similar to NETHIGH and can be found in Supplementary Figs. S1 and S2, respectively). Results were qualitatively similar to an alternative choice of contact-based EEI definition and survival p-value cutoff (Supplementary Figs. S3 and S4, respectively). Each point in the figure represents the mean number of edges per patient averaged over all patients across all 15 cancer types, and the vertical bars over the points represent the corresponding standard deviations.

Because the CPM value of 0.5 resulted in the maximum number of CRPEs, we chose it as the threshold to decide whether an exon is expressed or not for the rest of our analyses. An edge from the global EEIN was considered a CRPE if its presence or absence was significantly associated with patient survival in any of the 15 cancer types. Note that survival analysis was performed for each cancer type separately (Section 2.6). The total of 3,316, 9,224, and 13,030 CRPEs were derived from NETHIGH, NETMEDIUM, and NETLOW, respectively. Note that, because we identified an edge to be a CRPE if its expression was significantly associated with patient survival in at least one cancer type, the collection of CRPEs (e.g., the 3,316 edges corresponding to NETHIGH) might contain one or more edges that were not perturbed in one or more cancer types. Hence, for a given cancer type, only some of the CRPEs might be relevant. In the subsequent sections, we present the analysis of CRPEs in individual cancer types. Unless noted otherwise, all results presented in the main paper correspond to the CRPEs in NETHIGH, while the corresponding results (if applicable) for NETMEDIUM and NETLOW can be found in the Supplementary Materials.

### Cancer-relevant perturbed edges in individual cancer types

The number of CRPEs exhibited substantial variation across cancer types, from *∼*100 in PRAD to as many as *∼*1,700 in KIRC (Figure 5A and Supplementary Tables S1A-C). Many CRPEs were perturbed in multiple patients and the median number of patients in which a CRPE was perturbed varied from as high as 30 in BRCA to four in ESCA (Figure 5B). The number of CRPEs did not demonstrate significant correlation with the number of edges in control and condition samples, the total number of perturbed edges, and the number of patient samples used for the survival analysis (Supplementary Fig. S5), suggesting the absence of any systematic bias. At the same time, the number of CRPEs was significantly correlated with the number of genes (*p*-value *∼* 0.03; Figure 5C) that showed significant associations with patient survival based on their expression levels according to the study of Smith and Sheltzer (8) (Section 2.7). Note that in the latter study KIRC also stood out as the cancer type with the largest number of gene-survival associations. KIRC was also reported to have the highest proportion of cancer hallmark genes significantly associated with overall survival by Nagy et al. (5).

**Fig. 5:**
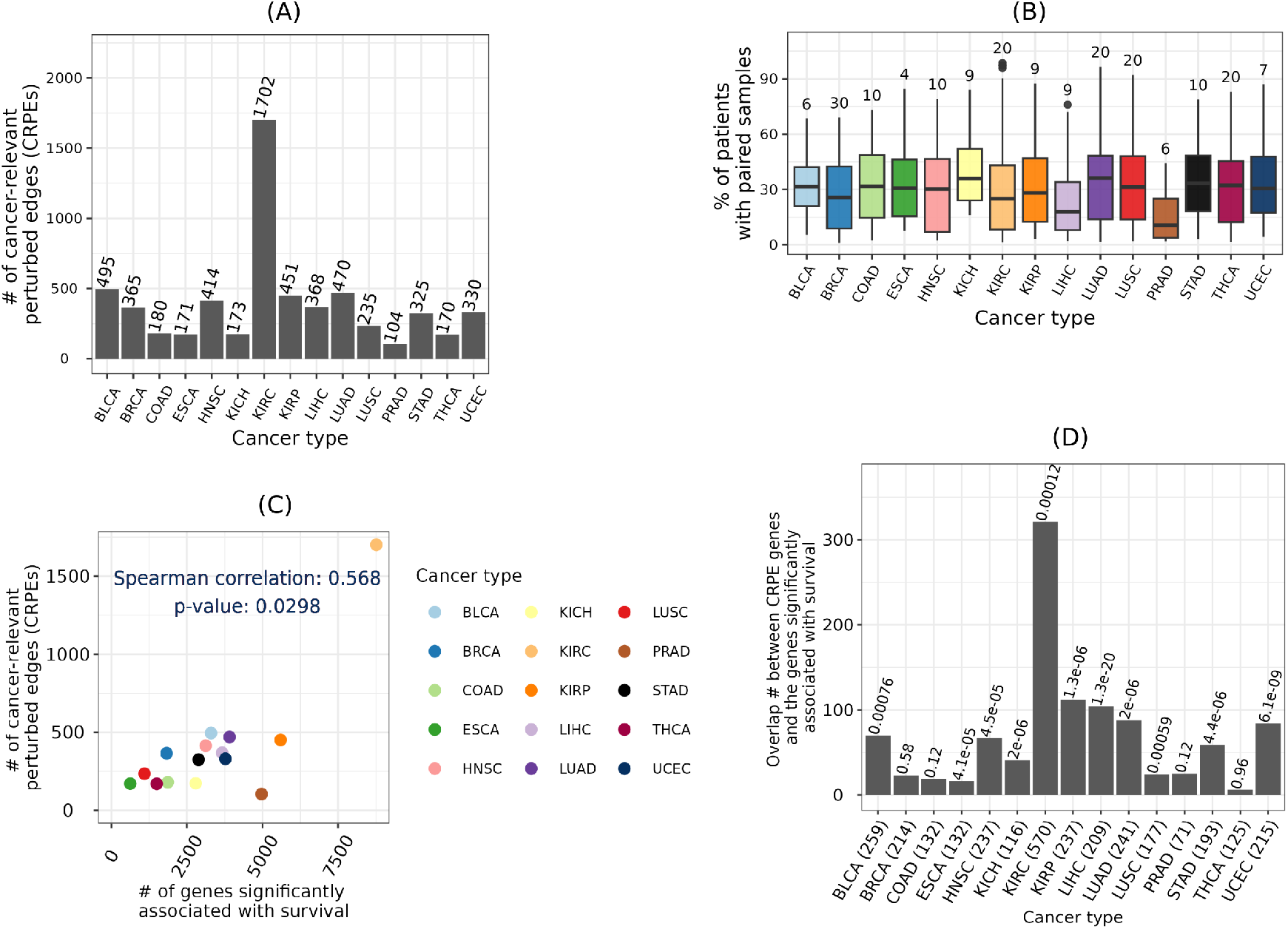
Cancer-relevant perturbed edges (CRPEs). (A) Counts of CRPEs associated with each cancer type. (B) Percentage of patients in which CRPEs were perturbed, with the median number of patients indicated on top of each box. (C) Correlation between the number of CRPEs and the number of genes significantly associated with patient survival according to Smith and Sheltzer (8). (D) Overlap between the genes from which CRPEs were derived and the genes significantly associated with patient survival according to Smith and Sheltzer (8). Number of genes from which the CRPEs were derived are shown within brackets along the x axis and the *q*-value of the overlap are shown on top of the bars. Results shown are for NETHIGH; results for NETMEDIUM and NETLOW are qualitatively similar (Supplementary Figs. S6 and S7).

Furthermore, for each cancer type we employed the hypergeometric test (Section 2.7) to evaluate whether the genes from which CRPEs were derived in this study show a statistically significant overlap with the genes associated with patient survival according to the study of Smith and Sheltzer (8). *p*-values of such overlaps across 15 cancer types were corrected using the Benjamini-Hochberg procedure to obtain the corresponding *q*-values. Significant (*q*-value< 0.001) overlaps were found for 11 out of 15 cancer types (Figure 5D). Because CRPEs are survival associated EEIs located on PPI interfaces, the above result highlights that a significant number of PPIs corresponding to the survival associated genes are also survival associated. Thus, exploring CRPEs can pinpoint PPIs that are survival biomarkers or cancer drug targets.

### Cancer-relevant perturbed edges and alternative splicing

We compared the number of CRPEs with the number of cancer type-related AS events that showed significant associations with patient survival reported in the study of Zhang et al. (34) (Section 2.8). Although the number of CRPEs did not correlate significantly (*p*-value *∼* 0.9) with the total number of AS events (data not shown), it did show a highly significant (*p*-value *∼* 0) correlation with the number of AS events that were significantly associated with the overall patient survival (Figure 6A). Because CRPEs are EEIs whose presence or absence is associated with patient survival (Section 3.2), exploring CRPEs can highlight AS events affecting protein complex formation, potentially revealing the mechanisms underlying cancer progression. Initially, we examined protein complex interfaces that were partially perturbed, meaning that, given an interface, some of its parts were never perturbed in any patient, while some of its other parts were CRPEs. We found several CRPEs that were part of partially perturbed protein complex interfaces, with the number of such CRPEs varying between 11 in COAD and 275 in KIRC (Figure 6B, Supplementary Tables S2A-C). Many of such CRPEs were perturbed in multiple patients, implying their significance for the corresponding cancer type (Figure 6C). For example, a CRPE between exon 7 (Ensembl ID “ENSE00003604293”) of PRKAB1 (UniProt ID “Q9Y478”) and exon 9 (Ensembl ID “ENSE00001454729”) of PRKAA2 (uniProt ID “P54646”) was perturbed in 26 out of 59 (*∼*44%) THCA patients. PRKAB1 has seven exons and PRKAA2 has nine exons. The protein complex between PRKAB1 and PRKAA2 has 21 EEIs, but only one (highlighted above) was perturbed while the remaining 20 EEIs were never perturbed in any of the THCA patients. These results exemplify the ability of our computational framework to highlight AS-related perturbations of protein complex interfaces potentially relevant for cancer prognosis.

**Fig. 6:**
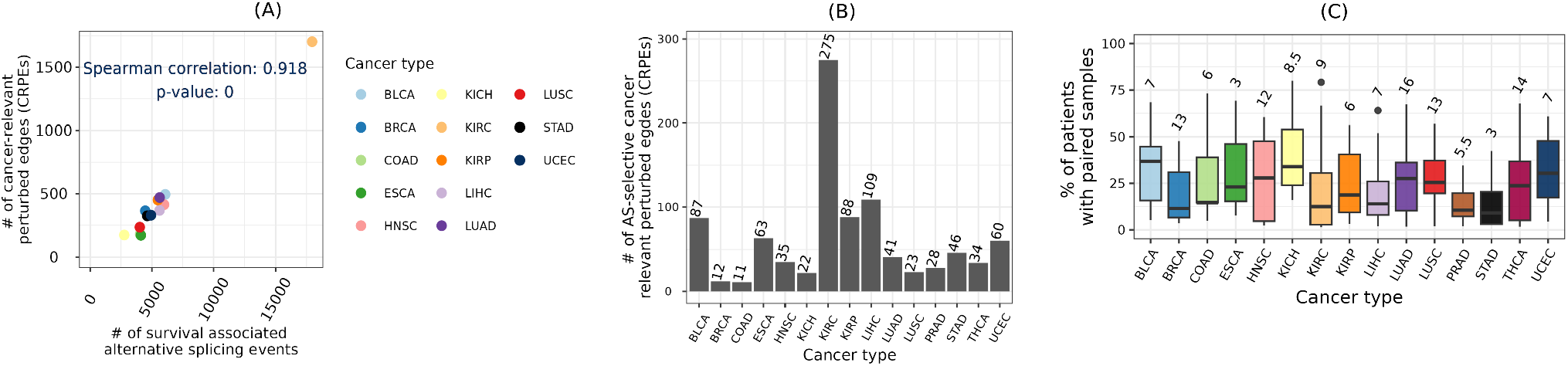
Cancer-relevant perturbed edges and alternative splicing (AS). (A) Correlation between the number of CRPEs and the number of AS events significantly associated with the overall patient survival. Data on AS events significantly associated with overall patient survival across all cancers except PRAD and THCA were obtained from the Supplementary Table S1 of Zhang et al. (34). (B) Number of CRPEs belonging to partially perturbed protein complexes (Supplementary Table S2). (C) Percentage of patients in which the CRPEs belonging to partially perturbed protein complexes were found, with the median number of patients indicated on top of each box. Results shown are for NETHIGH; results for NETMEDIUM and NETLOW are qualitatively similar (Supplementary Figs. S8 and S9).

### Shared and exclusive cancer-relevant perturbed edges

We found that CRPEs do not tend to be shared between different cancer types. Out of all 3,316 CRPEs in NETHIGH, 1,698 (i.e., *∼*51%) were unique to a given cancer type (Figure 7A). The largest (718) and smallest (11) counts of unique CRPEs were found in KIRC and PRAD, respectively. Out of 105 possible pairwise comparisons between 15 cancer types, only nine cancer type pairs had a significantly larger overlap in terms of the identified CRPEs than would be expected by chance, while 23 cancer type pairs had a significantly smaller overlap (Figure 7B). The largest degree of overlap, with three other cancer types, was exhibited by THCA and KIRP. The 170 CRPEs of THCA showed significantly more overlap with CRPEs of BRCA, KIRP, and STAD. The 452 CRPEs of KIRP showed significantly more overlap with CRPEs of THCA, PRAD, and LIHC. BRCA showed the smallest degree of overlap with other cancer types. Overall, these results imply that most of the CRPEs are cancer type specific and that different sets of biomolecular processes are associated with patient survival in different cancer types.

**Fig. 7:**
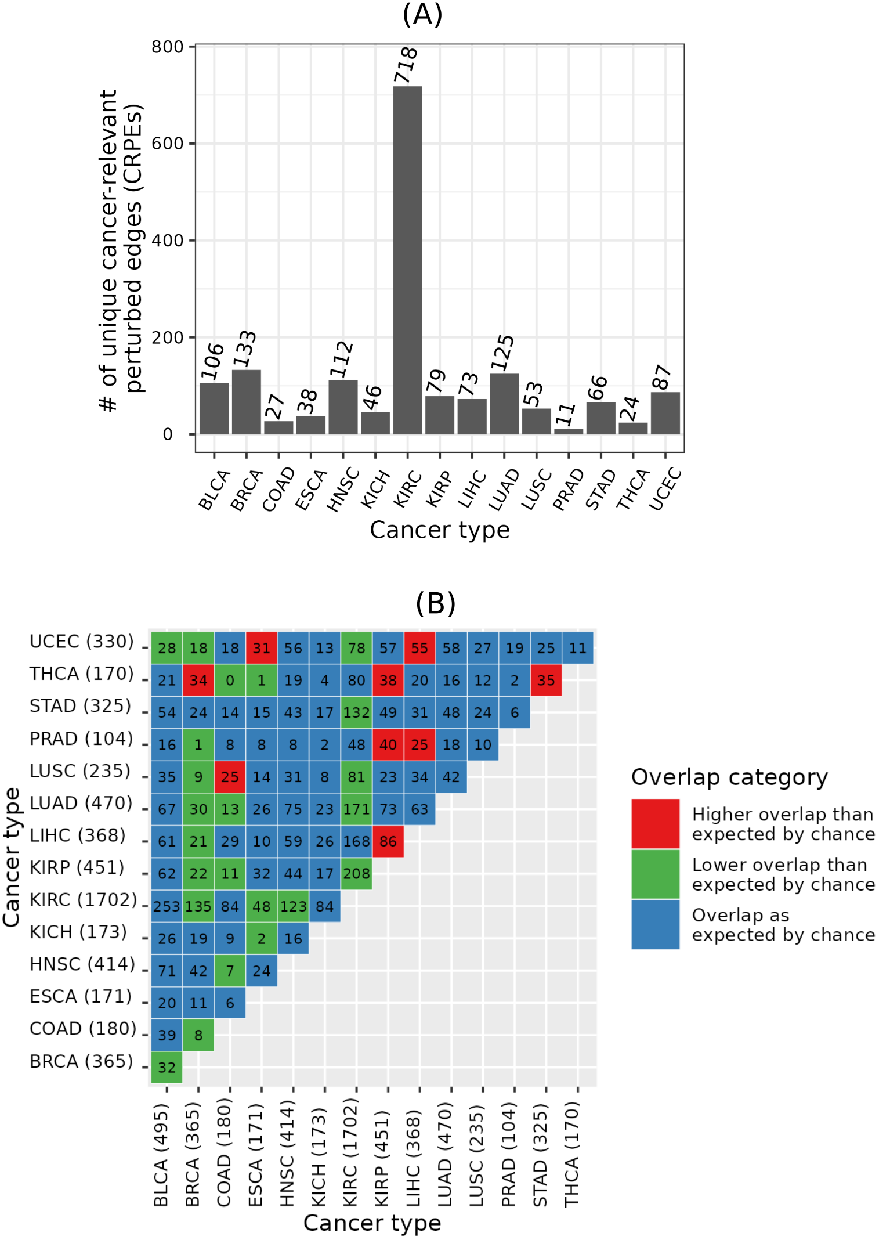
Unique and shared CRPEs across cancer types. (A) Counts of unique CRPEs in each cancer type. (B) Counts of CRPEs shared between cancer types. Given two cancer types, *p*-values of the overlap between their CRPEs with respect to the background set of edges in a global EEIN (here, NETHIGH) were calculated using the hypergeometric test and corrected using the Benjamini-Hochberg procedure to obtain the corresponding *q*-values. *q*-values less than or equal to 0.05 were considered significant. Numbers in each cell denote the counts of overlapping CRPEs between the corresponding cancer types, while the color indicates whether the overlap was significantly greater (red color) or smaller (green color) than expected by chance. For example, in the bottom-left green color cell, 32 CRPEs were shared between 495 CRPEs of BLCA and 365 CRPEs of BRCA, which was significantly less than would be expected by chance. Results shown are for NETHIGH; results for NETMEDIUM and NETLOW are qualitatively similar (Supplementary Figs. S10 and S11).

### Shared and exclusive metabolic pathways associated with cancer-relevant perturbed edges

To investigate which metabolic pathways were captured by CRPEs, we performed KEGG pathway enrichment analysis as follows. Given all exons involved in CRPEs of a cancer type, we evaluated whether any of the 285 KEGG pathways annotated within NETHIGH were enriched (FDR ≤ 0.05) using the hypergeometric test (Section 2.9). We did this for each of the 15 cancer types separately and obtained the corresponding cancer type-related enriched KEGG pathways. Note that most protein pairs among the 1,190 proteins in NETHIGH (Section 2.4) show pairwise sequence identities of less than 50% (Supplementary Fig. S12), which indicates that the NETHIGH proteins potentially come from diverse families and thus the corresponding 285 KEGG pathways represent a diverse set of metabolic pathways.

Out of all 219 KEGG pathways enriched in any cancer type, 95 (i.e., *∼*43%) were unique to a given cancer type (Figure 8A and Supplementary Tables S3A-C). The largest (36) and the smallest (zero) numbers of the uniquely enriched pathways were associated with BRCA and PRAD, respectively. Interestingly, while KIRC had the largest number of unique CRPEs (Figure 7A), it only had seven uniquely enriched pathways (Figure 8A). This implies that the unique KIRC CRPEs are predominantly enriched in those pathways that are also enriched in at least one other cancer type. Nevertheless, KIRC is uniquely enriched in the Sphingolipid signaling pathway (with a *q*-value of < 0.0003) that has been related to KIRC progression (38; 39) (Supplementary Fig. S15). Out of 105 possible pairs between the 15 cancer types, 16 pairs shared more enriched KEGG pathways than would be expected by chance. In particular, the 68 enriched pathways of STAD, as well as the 29 enriched pathways of LUSC, significantly overlapped with four other cancer types, while the 78 enriched pathways in BRCA showed no significant overlap with any other cancer type. Overall, these results imply that CRPEs in individual cancer types potentially cover different biological pathways.

**Fig. 8:**
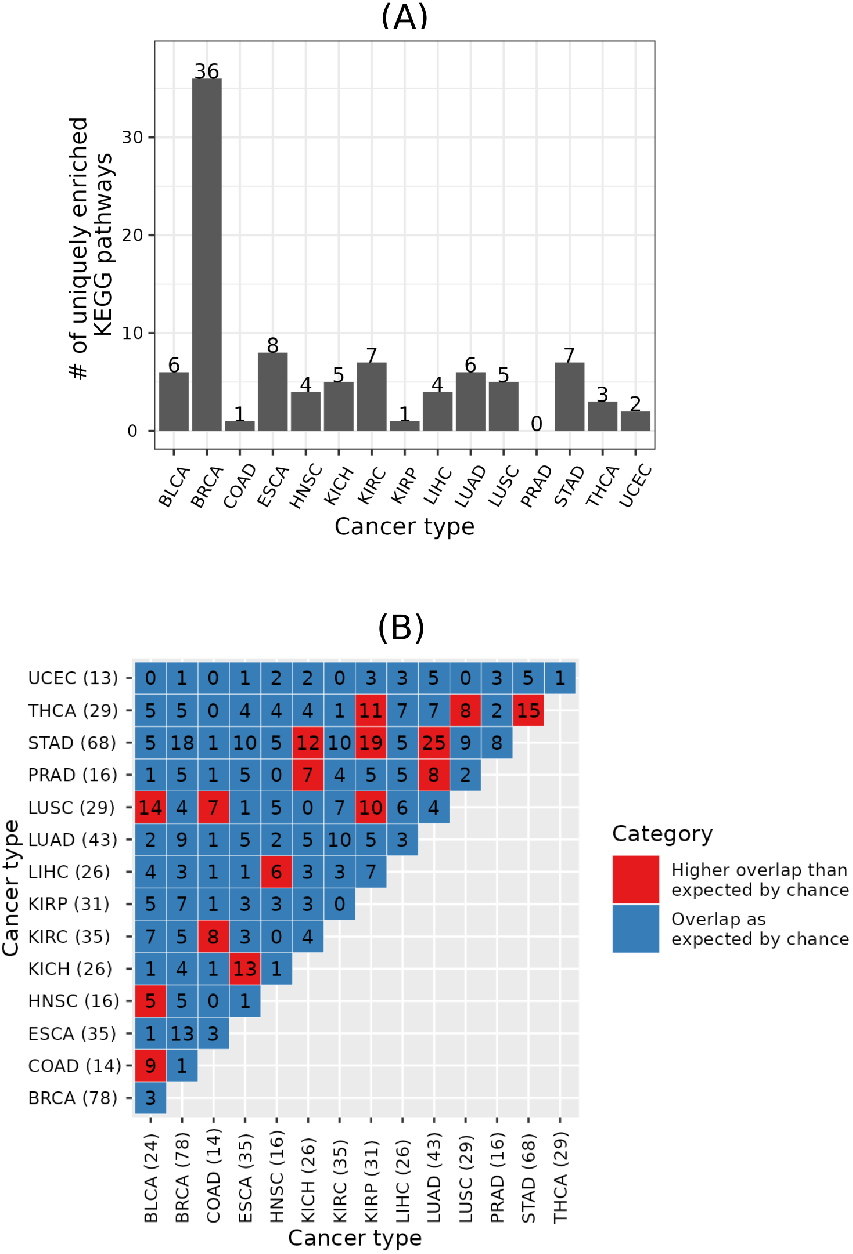
Unique and shared significantly enriched KEGG pathways across cancer types. (A) Number of uniquely enriched KEGG pathways in each cancer type. (B) Numbers of enriched KEGG pathways shared between cancer types. Given two cancer types, *q*-values of the overlap between their enriched KEGG pathways with respect to the total number of annotated KEGG pathways in a global EEIN (here, 285 KEGG pathways in NETHIGH) were computed, as for the overlapping CRPEs in Figure 7. Numbers in each cell denote the number of overlapping enriched KEGG pathways, while the color indicates whether the overlap was significantly higher (red color) than expected by chance. Results shown are for NETHIGH; results for NETMEDIUM and NETLOW are shown in Supplementary Figs. S13 and S14.

### Cancer-relevant perturbed edges resulting from different types of edge gain and loss events

We investigated the patterns of gain or loss events in cancer type-related CRPEs (Section 2.5). The proportion of CRPEs associated with different types of gain and loss events was similar across cancer types (Figure 9), with CRPEs gained in some patients and lost in other patients (“Gained or lost” category) being the most abundant category in all cancer types except LIHC. The next most abundant groups of CRPEs were associated with the “Strictly gained” and “Strictly lost” events, while “Patient-specific gained” and “Patient-specific lost” CRPEs were the least abundant. These results are expected given the heterogeneity of cancers, with only a few genomic features expected to be shared between patients of the same cancer type (40; 41; 42). Across different cancer types, the percentage of CRPEs associated with each type of gain and loss events varied greatly. For example, the representation of the “Gained or lost” category was lowest (*∼*40%) in LIHC and highest *∼*85% in THCA, while the “Strictly gained” and “Strictly lost” categories together were in the range between *∼*13% in THCA and *∼*55% in LIHC. Similarly, the percentage of CRPEs in the “Patient-specific gained” and “Patient-specific lost” categories together was lowest in KICH (0%) and highest (*∼*22%) in ESCA. Overall, these results imply the differences in the extent of heterogeneity among cancer types in terms of CRPEs.

**Fig. 9:**
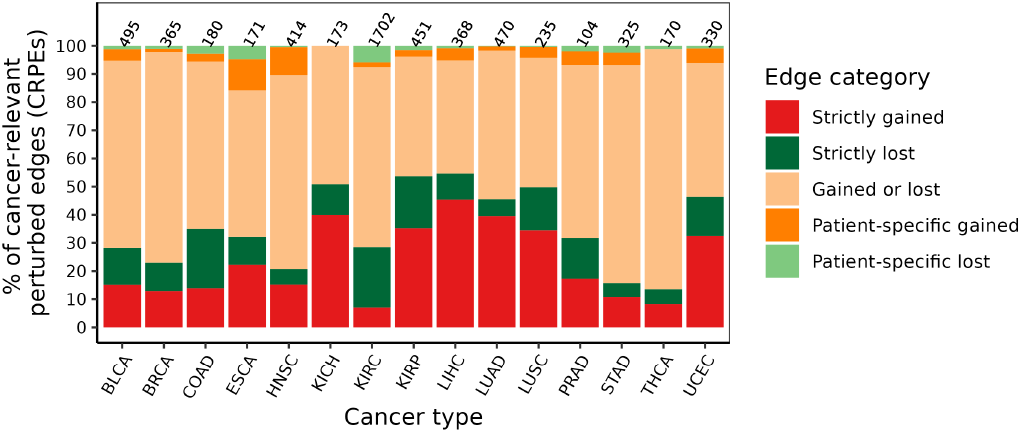
Distribution of cancer-relevant perturbed edge categories across cancer types. Results shown are for NETHIGH; results for NETMEDIUM and NETLOW are qualitatively similar (Supplementary Figs. S16 and S17).

### Cancer relevant perturbed edges as potential survival biomarkers

To investigate whether CRPEs can potentially be used as survival biomarkers, we evaluated whether CRPEs had the same statistical power (*p*-value) to be associated with patient survival using randomized patient survival data as with the real patient survival data (Section 2.10). Out of all 3,316 CRPEs from NETHIGH, we found 274 (*∼*8%) potential cancer survival biomarker CRPEs (BioCRPEs) (Supplementary Table S4). There was a large variation in BioCRPE occurrence across cancer types, with no BioCRPEs associated with KICH and the highest number of BioCRPEs (70) associated with KIRC (Figure 10A). Similar results were found by recent studies based on mRNA and protein expression data (3; 6), where significantly more mRNAs and proteins were identified as potential survival biomarkers in KIRC than in other cancer types. The number of BioCRPEs associated with higher (favorable) and lower (unfavorable) survival rates of patients varied across cancer types, with BRCA, COAD, ESCA, KIRC, and STAD having a higher number of favorable than unfavorable BioCRPEs, while for the rest of the cancer types we found the opposite trend (Figure 10A).

**Fig. 10:**
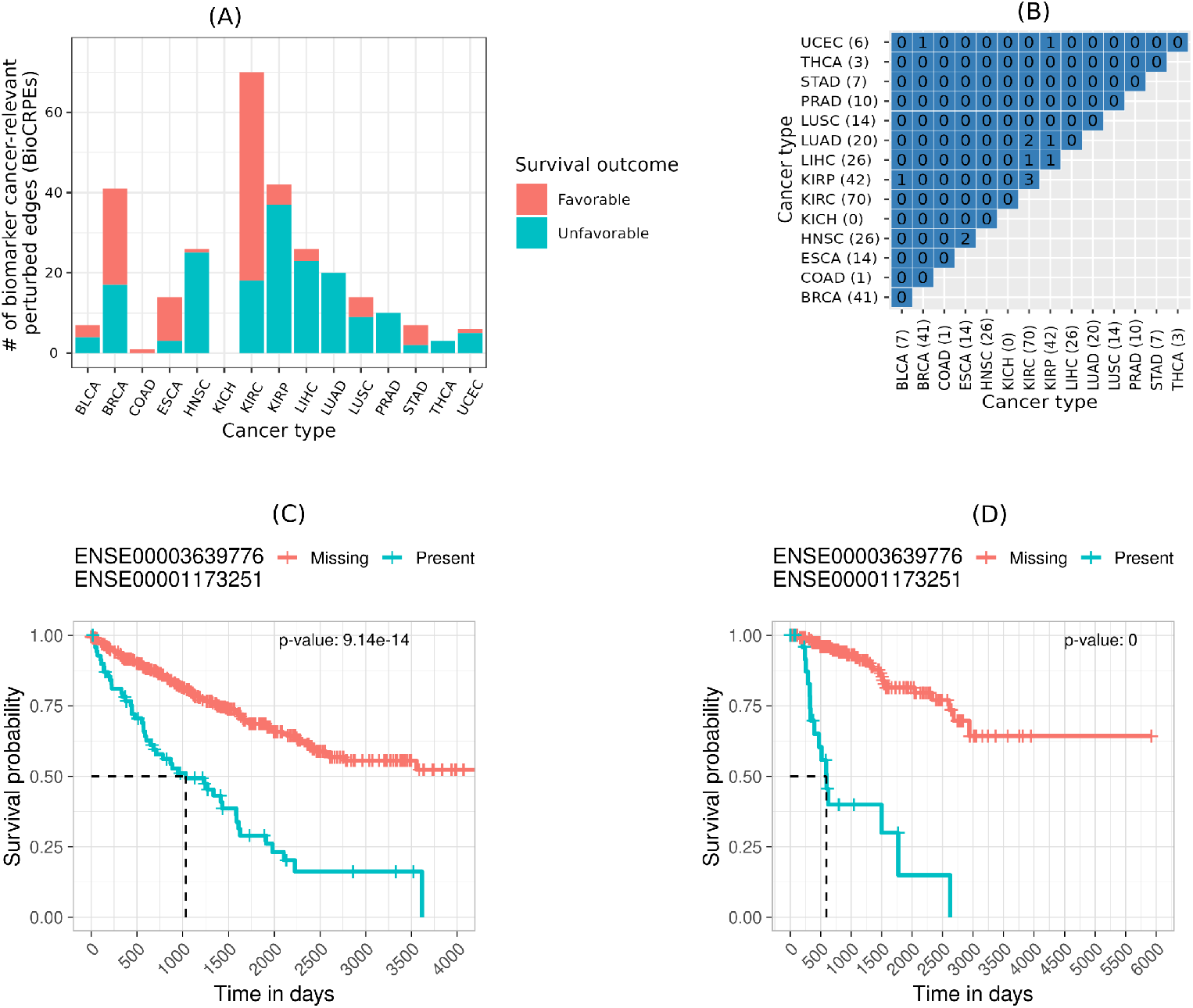
BioCRPEs across cancer types. (A) Number of BioCRPEs in each cancer type. (B) Pairwise overlaps between cancer types in terms of BioCRPEs. Given two cancer types, *q*-values of the overlap between their BioCRPEs were computed as for the overlaps between CRPEs in Figure 7. Results shown are for NETHIGH; results for NETMEDIUM and NETLOW are qualitatively similar (Supplementary Figs. S18 and S19). Results were qualitatively similar to an alternative choice of contact-based EEI definition and survival p-value cutoff (Supplementary Figs. S20 and S21, respectively). (C) Kaplan-Meier curve for a KIRC BioCRPE with the lowest log-rank *p*-value (displayed in the top-right area). Ensembl IDs of the participating exons are shown above the figure. (D) Kaplan-Meier curve for a KIRP BioCRPE with the lowest log-rank *p*-value.

We did not find significant overlaps between cancer types in terms of BioCRPEs (Figure 10B). Out of all 105 possible pairwise overlaps between BioCRPEs of the 15 cancer types, only nine were non-zero and none of them were significant, with the greatest number of BioCRPEs (three) shared between KIRC and KIRP. These results imply the different cancer types have almost a unique set of BioCRPEs. Interestingly, the most significantly associated unfavorable BioCRPE in both KIRC and KIRP was the same (Figure 10C-D), i.e., the interaction between exons with Ensembl IDs “ENSE00003639776” and “ENSE00001173251” that come from genes CCNB1 (UniProt ID “P14635”) and CDK1 (UniProt ID “P06493”), respectively. Additionally, this BioCRPE was absent from any other cancer type, implying that it is specific for the KIRC and KIRP groups of cancer types. Expression of CDK1 and CCNB1 genes has been shown to be correlated in cancer and higher expression of each of these genes has been reported to be a prognostic biomarker for KIRC and KIRP progression (43; 44). Our findings thus agree with the existing knowledge and hence our study has potential to unravel novel cancer survival biomarkers.

### Novel cancer biomarkers derived from perturbed edges

We investigated whether the identified BioCRPEs overlap with known cancer survival biomarkers. To do this, given a BioCRPE, we evaluated whether any of its two participating genes were significantly associated with survival according to the study of Smith and Sheltzer (8) (Section 2.7). We found that for many BioCRPEs, either one or both participating genes were associated with survival (Figure 11 and Supplementary Table S4). For example, out of the 41 BioCRPEs in BRCA, for 13 (*∼*32%) BioCRPEs at least one of the participating genes was a known survival biomarker, while for the rest of the 28 (*∼*68%) BioCRPEs none of the involved genes were known survival biomarkers. Thus, our study captures not only known but also novel survival biomarkers. In the following paragraph, as an example of how our findings can be further analyzed to gain novel insights into the importance of protein complexes in cancer, we present a specific example of two novel BioCRPEs related to BRCA.

**Fig. 11:**
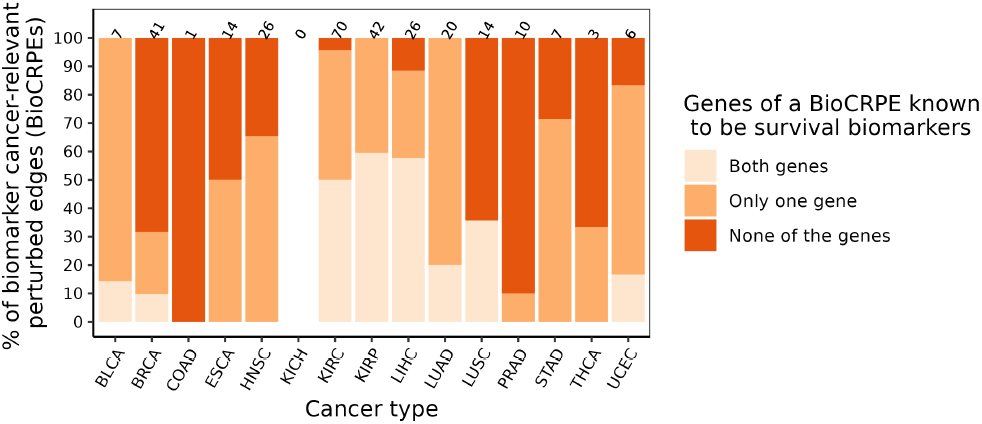
Number of novel BioCRPEs. For each cancer type, given the number of BioCRPEs (written on top of the corresponding bar), the percentage of known or novel BioCRPEs are outlined in different shades of orange.

We found two BioCRPEs that form an interface between subunits 2 (UniProt ID “Q9Y248”) and 4 (UniProt ID “Q9BRT9”) of the go-ichi-ni-san (GINS) protein complex. One BioCRPE is between exons with Ensembl IDs “ENSE00003585396” and “ENSE00001303150” from proteins Q9BRT9 and Q9Y248, respectively. The other BioCRPE is between exons with Ensembl IDs “ENSE00003554215” and “ENSE00001303150” from proteins Q9BRT9 and Q9Y248, respectively. The GINS complex has four protein subunits and helps in initiation and progression of DNA replication (45). Higher expression of the GINS subunit 2 has been related to poor prognosis in breast cancer patients (46), which is in line with our findings (Figure 12A). Looking closely into the protein-protein interaction interfaces between the subunits of the GINS complex (PDB ID 2E9X), we found that the interface between subunits 2 and 4 is stabilized by eight salt bridges and 20 hydrogen bonds (interface information taken from the PISA server using PDB ID 2E9X). Interestingly, parts of this interface overlap with the exons involved in the above two BioCRPEs, capturing the majority (five out of eight) salt bridges and some (four out of 20) hydrogen bonds of the interface (Figure 12B). This may imply that the interactions between the exons captured by the above two BioCRPEs are important for the stability of the corresponding interface in the GINS complex.

**Fig. 12:**
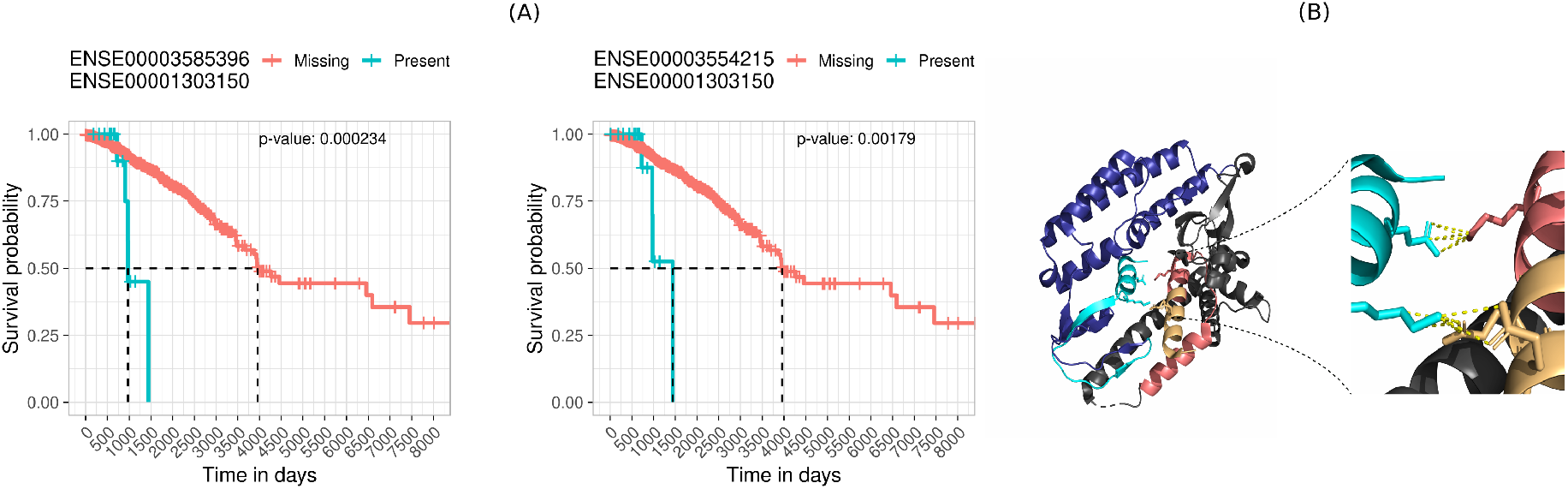
An example of two novel BioCRPEs related to BRCA. (A) Kaplan-Meier curves corresponding to the two BioCRPEs. (B) The left portion of the figure shows the 3D structure of the part of the GINS complex containing subunits 2 (Q9Y248) and 4 (Q9BRT9). The protein Q9Y248 is shown in blue color with the associated exon “ENSE00001303150” participating in the two BioCRPEs in cyan color. The protein Q9BRT9 is shown in gray color with the exons “ENSE00003554215” and “ENSE00003585396” participating in the two BioCRPEs in yellow and pink colors, respectively. The right portion of the figure highlights inter-protein interactions (within 4Å, shown as yellow dashed lines) between the exons participating in the two BioCRPEs.

## Conclusion

In this study, transcriptomics data were integrated with structurally resolved protein complexes to identify perturbed exon-exon interactions (EEIs) and to investigate their prognostic value for patient survival across 15 different cancer types. Because most alternative splicing (AS) events add or remove exons to produce alternative protein isoforms, EEIs indirectly capture the effect of AS on protein-protein interaction (PPI) interfaces. The number of cancer-relevant perturbed EEIs significantly correlates with the previously reported number of prognostic AS events. Because EEIs were derived from PPI interfaces, further exploration of such perturbed EEIs can identify prognostic AS events that are likely to affect PPIs. This information could be helpful in guiding the development of anti-cancer drugs that target specific PPIs affected by prognostic AS events.

Furthermore, we found many cancer-relevant perturbed EEIs that are uniquely present in individual cancer types and capture different molecular processes in different cancer types. Further exploration of cancer type-related perturbed EEIs could potentially shed light on the molecular mechanisms driving cancer progression. We identified many novel prognostic EEIs involving interacting proteins for which no association with patient survival has been previously reported. These proteins constitute valuable candidates for experimental validation to further elucidate their functional roles in cancer. Note that AS-related molecular changes have been found to be similar in tumors and their corresponding microenvironments (47; 48). Because EEI perturbations in this study were captured by comparing tumor samples with the paired healthy samples, the proximity of healthy samples with the tumor microenvironment could negatively affect the extent of the observed EEI perturbations. Future work in this direction could uncover valuable insights.

Modeling interactions between exons as a network provides a framework where one can directly map the exon expressions to study AS-related PPI perturbations. As the number of structurally resolved (or accurately modeled) protein complexes grows, the coverage of EEI networks is poised to increase, further improving the usefulness of our computational framework for exploring the impact of AS-related PPI network perturbations on clinical outcomes in cancer.

## Data availability

RNA-seq data were downloaded from the The Cancer Genome Atlas. Three-dimensional protein structural data were downloaded from the Protein Data Bank. Data to reproduce results is available at: https://doi.org/10.5281/zenodo.12917451. Code to reproduce results are present in the GitHub repository: https://github.com/KhaliqueN/Cancer_prognostic_EEIs.

## Author contributions statement

Conceived the study: K.N. and D.F. Collected and processed the data: K.N. Designed the experiments: K.N. and D.F. Performed the experiments: K.N. Analysed the results: K.N. and D.F. Wrote the paper: K.N., and D.F. Read and approved the paper: K.N., C.S., K.W., J.B. and D.F. Supervised the study: D.F.

## Acknowledgments

This work was supported by Universität Hamburg and HamburgX grant LFF-HHX-03 to the Center for Data and Computing in Natural Sciences (CDCS) from the Hamburg Ministry of Science, Research, Equalities and Districts. This project is funded by the European Union under grant agreement No. 101057619. Views and opinions expressed are however those of the author(s) only and do not necessarily reflect those of the European Union or European Health and Digital Executive Agency (HADEA). Neither the European Union nor the granting authority can be held responsible for them. This work was developed as part of the ASPIRE project and is funded by the German Federal Ministry of Education and Research (BMBF) under grant number 031L0287B.

