## Supplementary materials for "Prognostic importance of splicing-triggered aberrations of protein complex interfaces in cancer"

### I Supplementary Sections

### S1. Key results of the study using an alternative contact-based EEI definition

In Section 2.3 of the main paper, using the contact-based approach, we defined two exons to interact if they showed at least one residue-residue contact. Here, we repeated our study using an alternative contact-based EEI definition, where we defined two exons to interact if they showed at least five residue-residue contacts. This approach resulted in 6,617 contact-based EEIs among 4,266 exons. We used this new contact-based EEIN, along with the energy-based and evolution-based EEINs (Section 2.3), to construct a new NETHIGH EEIN with 3,203 EEIs and 2,225 exons. We found that the key results (summarized below) corresponding to this new NETHIGH EEIN are qualitatively similar to the key results corresponding to the original NETHIGH EEIN that we presented in the main paper.

The CPM threshold of 0.5 for exon expression resulted in the maximum number (1,679) of CRPEs (Supplementary Fig. S3 A). The number of CRPEs did not demonstrate significant correlation with the number of edges in control and condition samples, the total number of perturbed edges, and the number of patient samples used for the survival analysis (Supplementary Figure S3 B), suggesting the absence of any systematic bias. Also, the number of CRPEs was significantly correlated with the number of genes (Supplementary Fig S3 C left) that showed significant associations with patient survival based on their expression levels according to Smith and Sheltzer [1]. The number of CRPEs showed a highly significant correlation with the number of alternative splicing events that were significantly associated with the overall patient survival in Zhang et al. [2] (Supplementary Fig S3 C right). We found that CRPEs or their enriched KEGG pathways do not tend to be shared between different cancer types (Supplementary Fig S20 A). We found that the proportion of CRPEs associated with different types of gain and loss events was similar across cancer types (Supplementary Fig S20 B). Finally, we found many cancer type-related BioCRPEs with non-significant overlaps across cancer types (Supplementary Fig S20 C).

### S2. Key results of the study using an alternative $p$-value cutoff for survival analysis

In Section 3.1 of the main paper, we identified CRPEs as those EEIs whose perturbations significantly correlated with survival in at least one cancer type using a survival analysis $p$-value cutoff of 0.05. The selected CRPEs were then used in the downstream analyses of the paper. Here, we repeated our study for the most confident EEIN (NETHIGH) using an alternative survival analysis $p$-value cutoff of 0.01. We found that the key results corresponding to the survival analysis $p$-value cutoff of 0.01 (summarized below) are qualitatively similar to the results corresponding to the survival analysis $p$-value cutoff of 0.05 that we presented in the main paper.

The CPM threshold of 0.5 for exon expression resulted in the maximum number (2,250) of CRPEs (Supplementary Fig. S4 A). The number of CRPEs did not demonstrate significant correlation with the number of edges in control and condition samples, the total number of perturbed edges, and the number of patient samples used for the survival analysis (Supplementary Figure S4 B), suggesting the absence of any systematic bias. Also, the number of CRPEs was significantly correlated with the number of genes (Supplementary Fig S4 C left) that showed significant associations with patient survival based on their expression levels according to Smith and Sheltzer [1]. The number of CRPEs showed a highly significant correlation with the number of alternative splicing events that were significantly associated with the overall patient survival in Zhang et al. [2] (Supplementary Fig S4 C right). We found that CRPEs or their enriched KEGG pathways do not tend to be shared between different cancer types (Supplementary Fig S21A). We found that the proportion of CRPEs associated with different types of gain and loss events was similar across cancer types (Supplementary Fig S21B). Finally, we found many cancer type-related BioCRPEs with non-significant overlaps across cancer types (Supplementary Fig S21C).

### S3. Dependence between the number of edges and the exon expression value threshold

Here, we complement Section 3.1 of the main paper by explaining the details about how the number of perturbed edges or cancer relevant perturbed edges (CRPEs) vary with different exon expression threshold values. Theoretically, for the two extreme expression threshold values of 0 and infinity, the number of perturbed edges would be zero because the control edges and the condition edges would include either all (for the threshold of 0) or none (for the threshold of infinity) of the edges in the entire exon-exon interaction network. Hence, if there is at least one expression threshold X between 0 and infinity with a non-zero number of perturbed edges, then we would expect to see the number of perturbed edges to first increase from threshold 0 to X and then decrease from X to infinity, which theoretically justifies the results that we observe in Figure 4 of the main paper and in Supplementary Figs S1 and S2. That is, although the numbers of edges in the paired control and condition samples decrease along the CPM values of 0.05, 0.1, 0.2, 0.5, 1, and 2 (as expected), the number of perturbed edges first increases for the expression values from 0.05 to 0.5 and then decreases for the expression values from 0.5 to 2.

### II Supplementary Figures

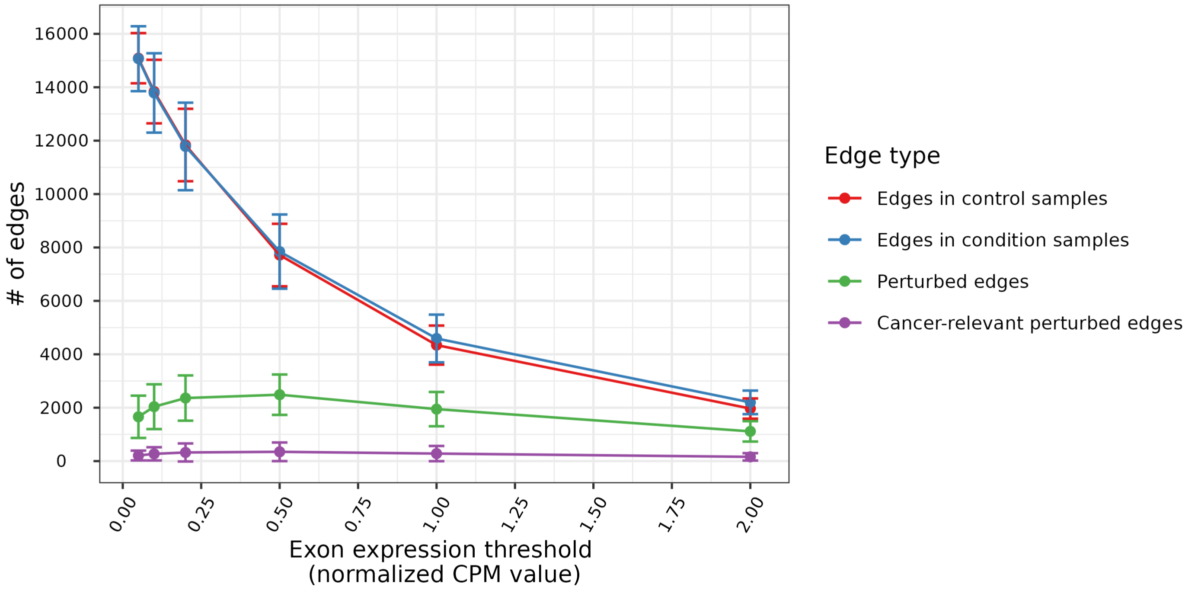


Supplementary Fig. S1: Dependence of the number of edges on the choice of expression value threshold (results shown for NETMEDIUM). Each point in the figure represents the mean number of edges per patient averaged over all patients across all 15 cancer types, and the vertical bars over the points represent the corresponding standard deviations.

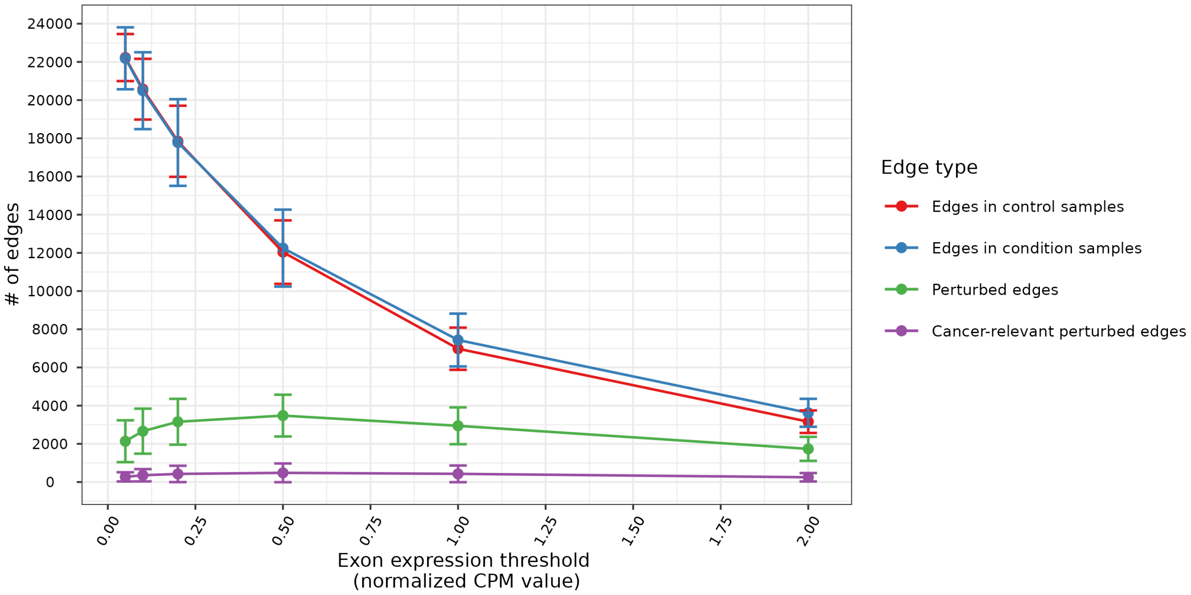


Supplementary Fig. S2: Dependence of the number of edges on the choice of exon expression value threshold (results shown for NETLOW). Each point in the figure represents the mean number of edges per patient averaged over all patients across all 15 cancer types, and the vertical bars over the points represent the corresponding standard deviations.

(A)

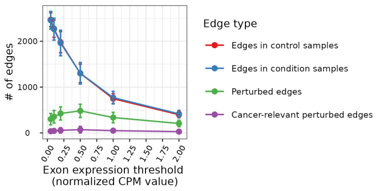


(B)

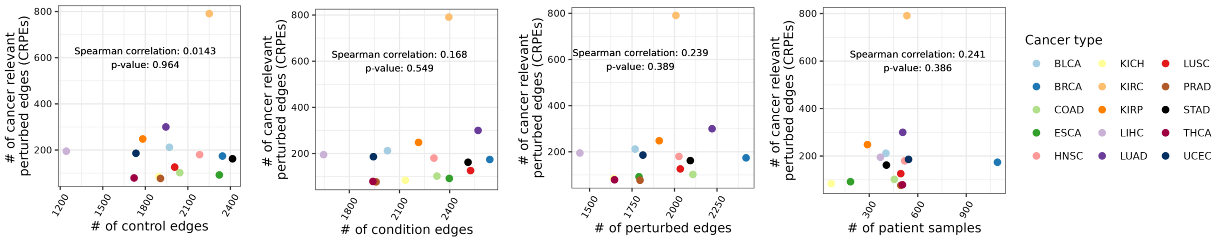


(C)

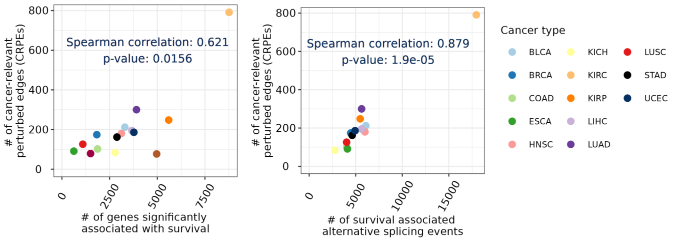


Supplementary Fig. S3: Key results corresponding to the Sections 3.1 to 3.3 of the main paper using an alternative definition of contact-based EEIs, where we defined a pair of exons to interact if and only if they have at least five residue-residue contacts as per our residue-residue definition in Section 2.3 of the main paper. All results are shown for NETHIGH created using the new contact-based EEI definition. (A) Dependence of the number of edges on the choice of expression value threshold. (B) Correlation between the number CRPEs and, from left to right, (1) the total number of edges in control samples, (2) the total number of edges in condition samples, (3) the total number of perturbed edges, and (4) the total number of patients with at least one condition sample and overall survival information. (C) Correlation between the number of CRPEs and, from left to right, (1) the number of genes significantly associated with patient survival according to Smith and Sheltzer [1] and (2) the number of AS events significantly associated with the overall patient survival Zhang et al. [2].

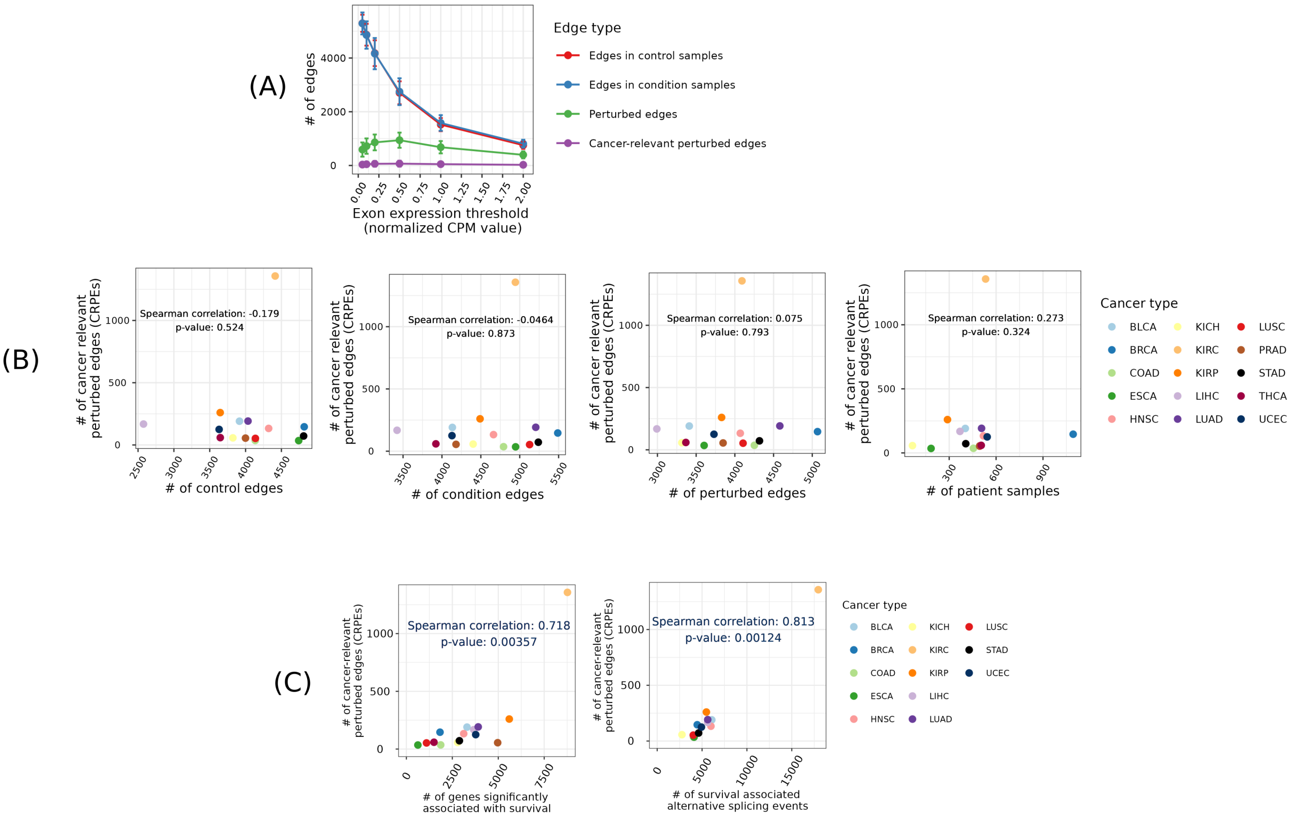


Supplementary Fig. S4: Key results corresponding to the Sections 3.1 to 3.3 of the main paper using an alternative $p$-value cutoff for survival analysis. All results are shown for NETHIGH. (A) Dependence of the number of edges on the choice of expression value threshold. (B) Correlation between the number CRPEs and, from left to right, (1) the total number of edges in control samples, (2) the total number of edges in condition samples, (3) the total number of perturbed edges, and (4) the total number of patients with at least one condition sample and overall survival information. (C) Correlation between the number of CRPEs and, from left to right, (1) the number of genes significantly associated with patient survival according to Smith and Sheltzer [1] and (2) the number of AS events significantly associated with the overall patient survival Zhang et al. [2].

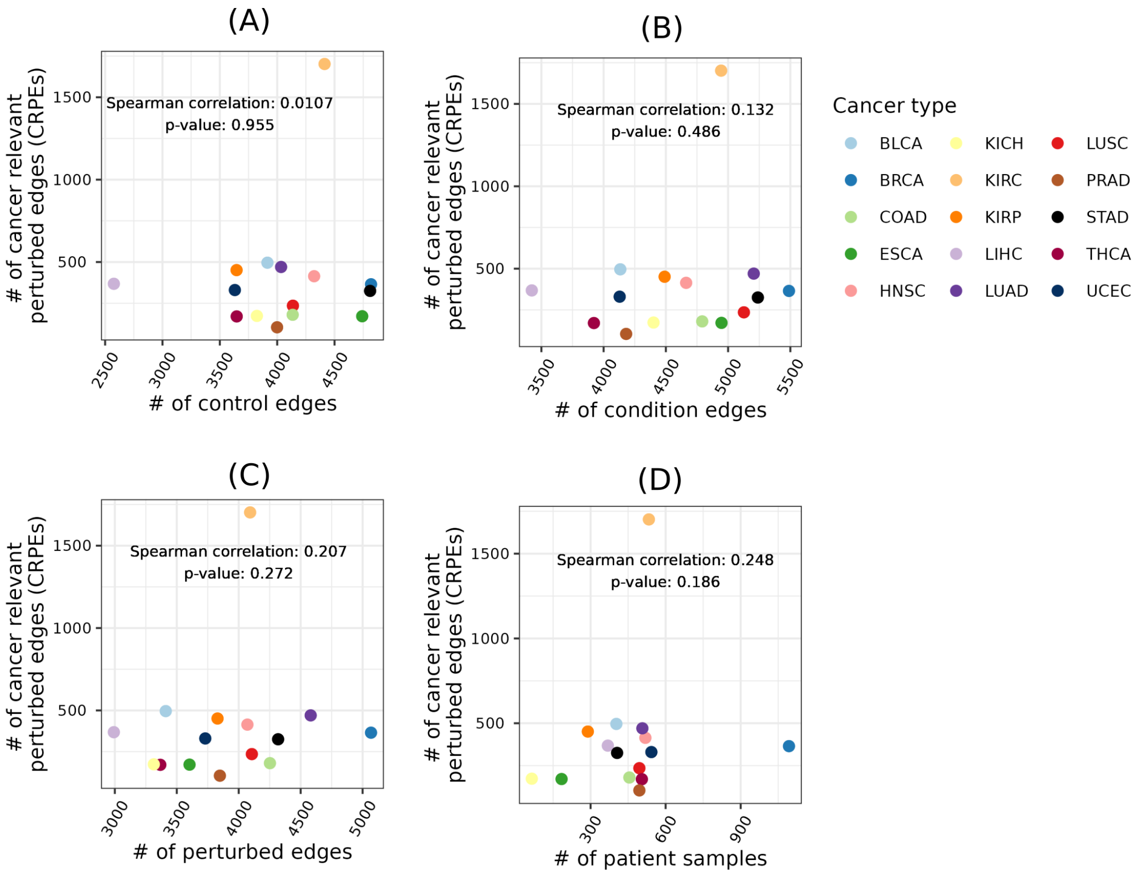


Supplementary Fig. S5: Correlation between the number cancer relevant perturbed edges (CRPEs) using NETHIGH and (A) the total number of edges in control samples, (B) the total number of edges in condition samples, (C) the total number of perturbed edges, and (D) the total number of patients with at least one condition sample and overall survival information.

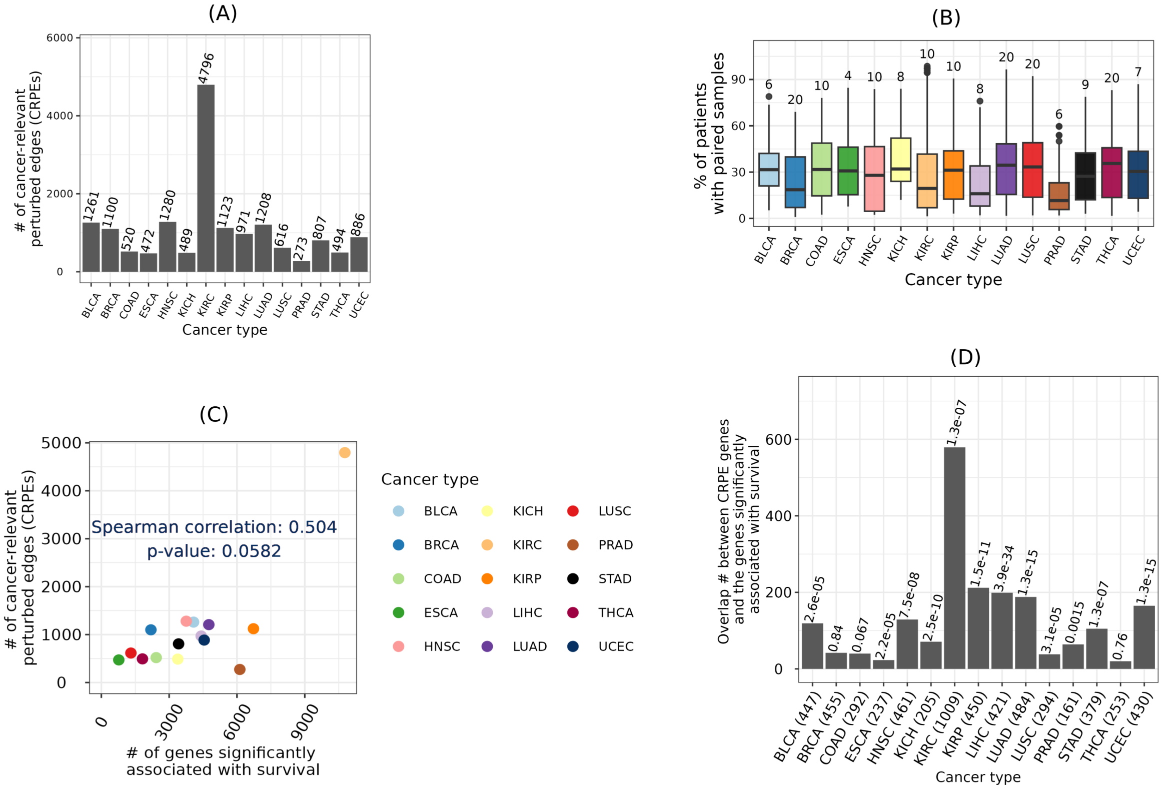


Supplementary Fig. S6: Cancer-relevant perturbed edges (CRPEs) vs genes with respect to NETMEDIUM. (A) Counts of CRPEs associated with each cancer type. (B) Percentage of patients in which the CRPEs were perturbed, with the median number of patients indicated on top of each box. (C) Correlation between the number of CRPEs and the number of genes significantly associated with patient survival according to [1]. (D) Overlap between the genes from which CRPEs were derived and the genes significantly associated with patient survival according to [1]. Number of genes from which the CRPEs were derived are shown within brackets along the x axis and the $q$-value of the overlap are shown on top of the bars.

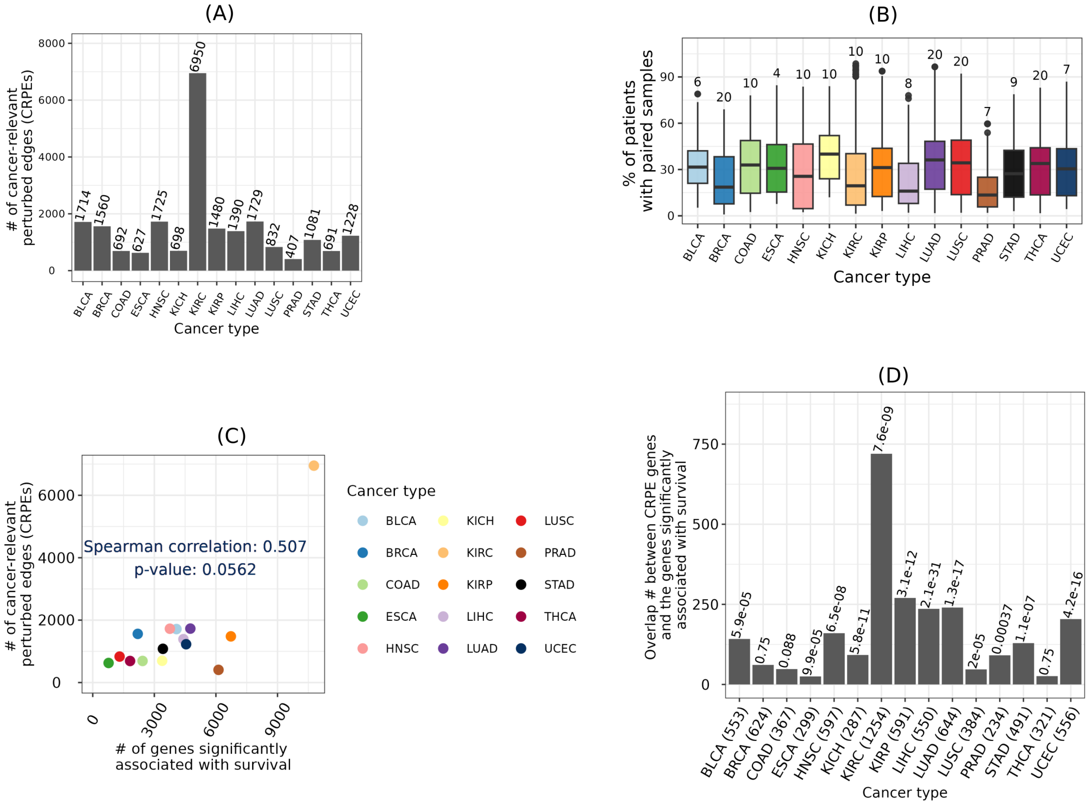

Supplementary Fig. S7: Cancer-relevant perturbed edges (CRPEs) vs genes with respect to NETLOW. (A) Counts of CRPEs associated with each cancer type. (B) Percentage of patients in which the CRPEs were perturbed, with the median number of patients indicated on top of each box. (C) Correlation between the number of CRPEs and the number of genes significantly associated with patient survival according to Smith and Sheltzer [1]. (D) Overlap between the genes from which CRPEs were derived and the genes significantly associated with patient survival according to [1]. Number of genes from which the CRPEs were derived are shown within brackets along the x axis and the $q$-value of the overlap are shown on top of the bars

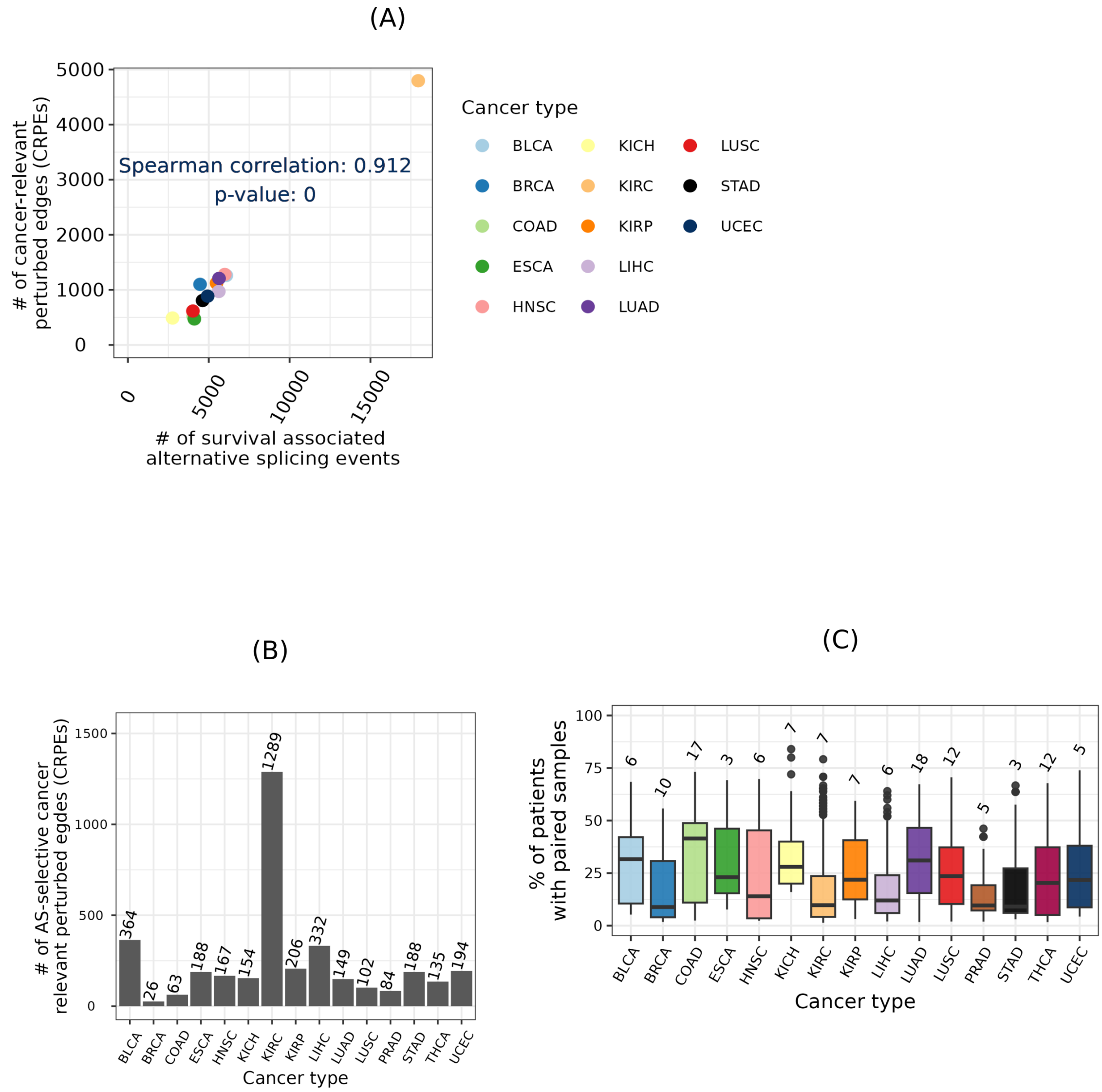


Supplementary Fig. S8: Cancer-relevant perturbed edges and alternative splicing (AS) with respect to NETMEDIUM. (A) Correlation between the number of CRPEs and the number of AS events significantly associated with the overall patient survival. Data on AS events significantly associated with overall patient survival across all cancers except PRAD and THCA were obtained from the Supplementary Table S1 of [2]. (B) Number of CRPEs belonging to partially perturbed protein complexes. (C) Percentage of patients in which the CRPEs belonging to partially perturbed protein complexes were found, with the median number of patients indicated on top of each box.

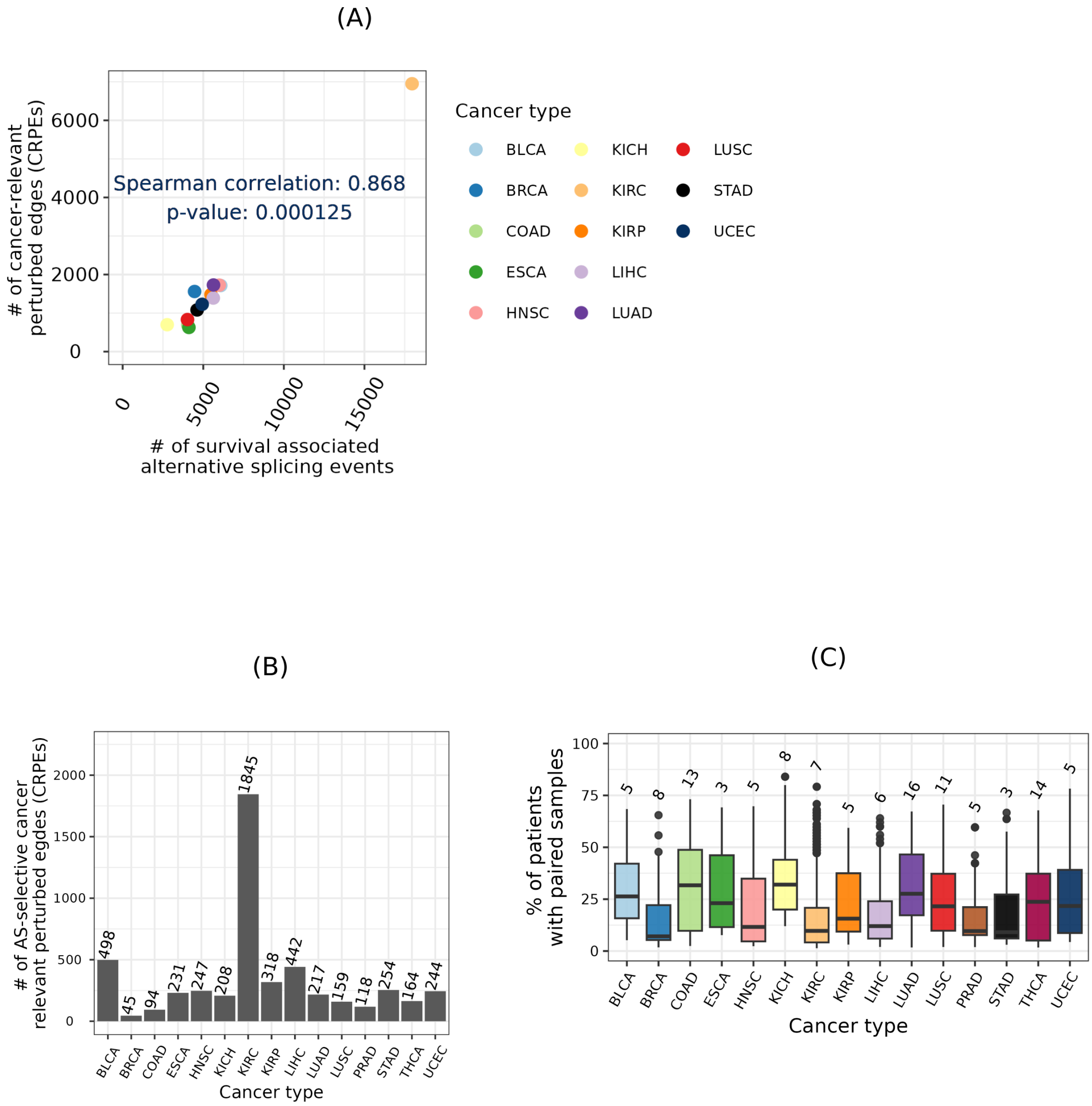


Supplementary Fig. S9: Cancer-relevant perturbed edges and alternative splicing (AS) with respect to NETLOW. (A) Correlation between the number of CRPEs and the number of AS events significantly associated with the overall patient survival. Data on AS events significantly associated with overall patient survival across all cancers except PRAD and THCA were obtained from the Supplementary Table S1 of [2]. (B) Number of CRPEs belonging to partially perturbed protein complexes. (C) Percentage of patients in which the CRPEs belonging to partially perturbed protein complexes were found, with the median number of patients indicated on top of each box.

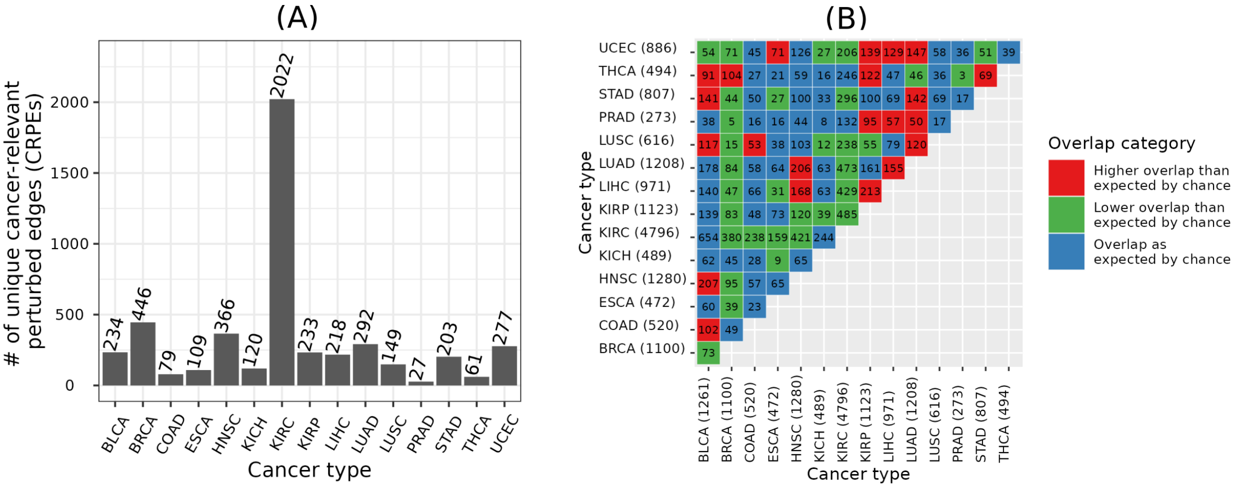

Supplementary Fig. S10: Unique and shared CRPEs across cancer types using NETMEDIUM. (A) Counts of unique CRPEs in each cancer type. (B) Counts of CRPEs shared between cancer types. Given two cancer types, $p$-values of the overlap between their CRPEs with respect to the background set of edges in a global EEIN were calculated using the hypergeometric test and corrected using the Benjamini-Hochberg procedure to obtain the corresponding q-values. q-values less than or equal to 0.05 were considered significant. Numbers in each cell denote the counts of overlapping CRPEs between the corresponding cancer types, while the color indicates whether the overlap was significantly greater (red color) or smaller (green color) than expected by chance.

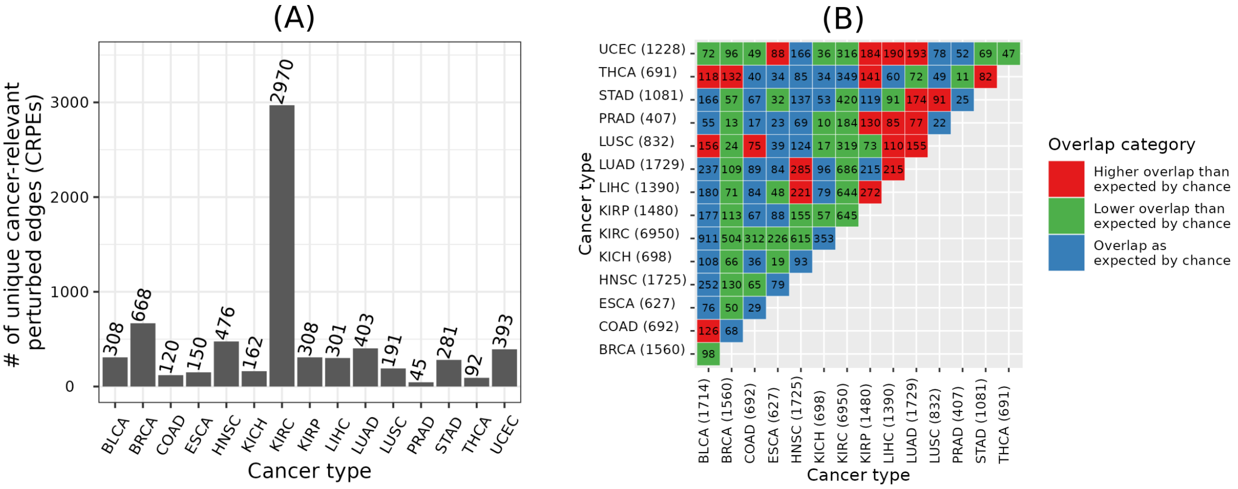

Supplementary Fig. S11: Unique and shared CRPEs across cancer types using NETLOW. (A) Counts of unique CRPEs in each cancer type. (B) Counts of CRPEs shared between cancer types. Given two cancer types, $p$-values of the overlap between their CRPEs with respect to the background set of edges in a global EEIN were calculated using the hypergeometric test and corrected using the Benjamini-Hochberg procedure to obtain the corresponding q-values. q-values less than or equal to 0.05 were considered significant. Numbers in each cell denote the counts of overlapping CRPEs between the corresponding cancer types, while the color indicates whether the overlap was significantly greater (red color) or smaller (green color) than expected by chance.

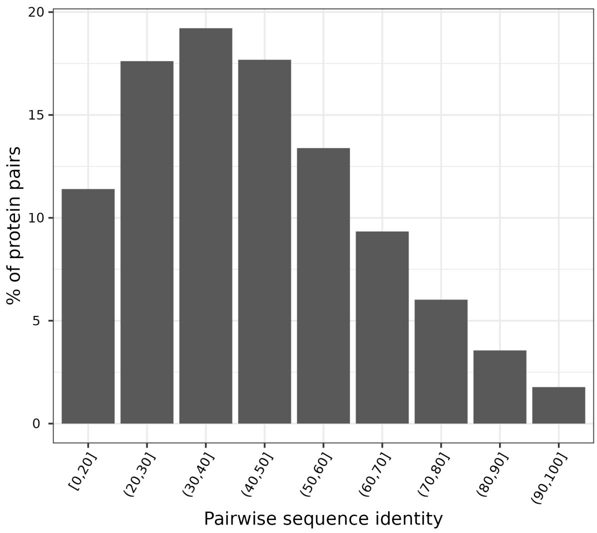


Supplementary Fig. S12: Pairwise sequence identities of proteins in NETHIGH. Given a pair of proteins, the sequence identities were measured as the number of matching amino acids in the corresponding global alignment divided by the total number of amino acids in the shorter protein sequence.

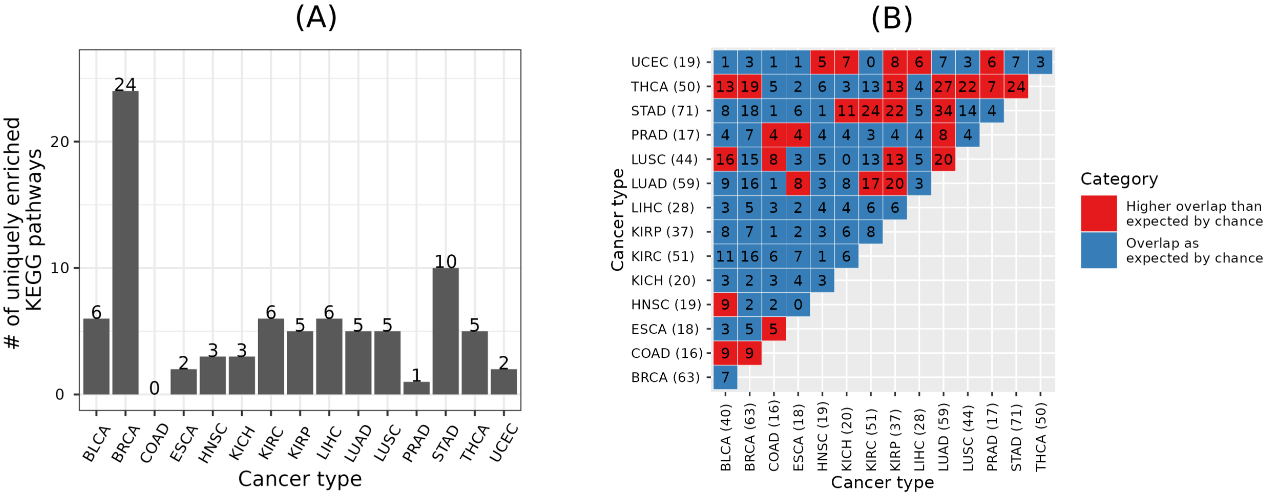


Supplementary Fig. S13: Unique and shared significantly enriched KEGG pathways across cancer types using NETMEDIUM. (A) Number of uniquely enriched KEGG pathways in each cancer type. (B) Numbers of enriched KEGG pathways shared between cancer types. Given two cancer types, q-values of the overlap between their enriched KEGG pathways were computed, as for the overlapping KEGG pathways in Figure 8. Numbers in each cell denote the number of overlapping enriched KEGG pathways, while the color indicates whether the overlap was significantly higher (red color) than expected by chance.

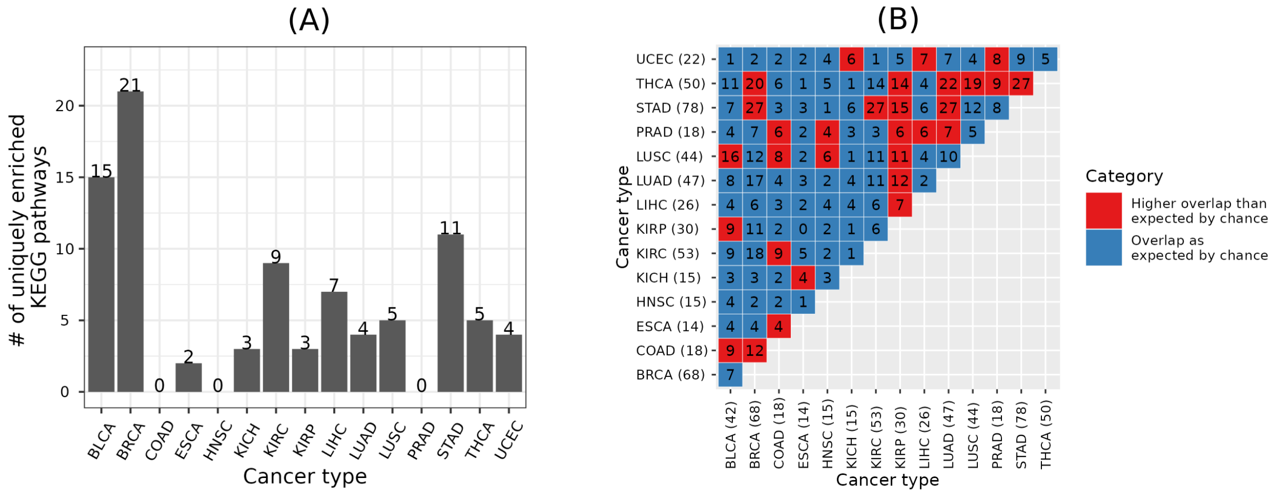


Supplementary Fig. S14: Unique and shared significantly enriched KEGG pathways across cancer types using NETLOW. (A) Number of uniquely enriched KEGG pathways in each cancer type. (B) Numbers of enriched KEGG pathways shared between cancer types. Given two cancer types, q-values of the overlap between their enriched KEGG pathways were computed, as for the overlapping KEGG pathways in Figure 8. Numbers in each cell denote the number of overlapping enriched KEGG pathways, while the color indicates whether the overlap was significantly higher (red color) than expected by chance.

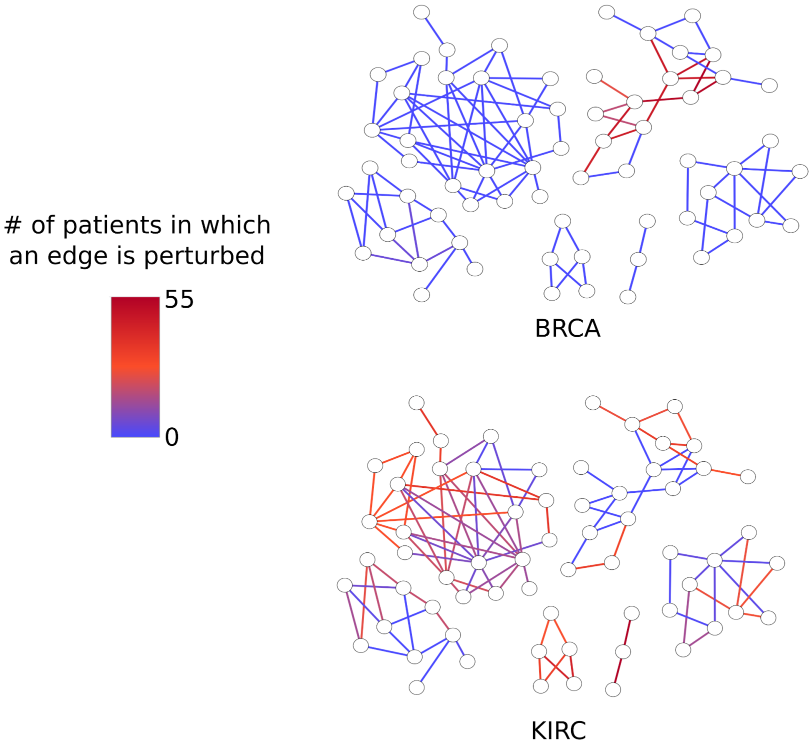


Supplementary Fig. S15: NETHIGH CRPE subnetwork (with 65 exons or nodes and 101 EEIs or edges) of the Sphingolipid signaling pathway (KEGG ID hsa04071). The edge color represents the number of patients in which the edge was detected as a CRPE. The results highlight the significant difference between EEI perturbations of KIRC in comparison to EEI perturbations of BRCA.

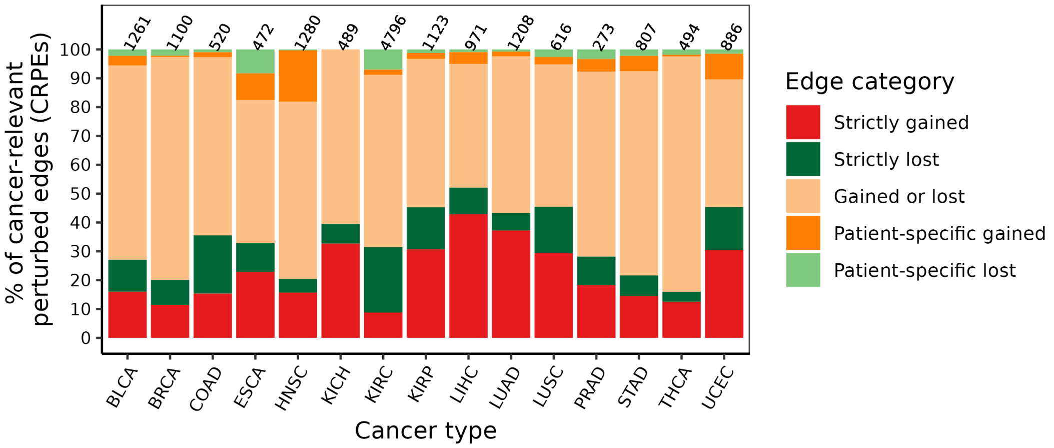


Supplementary Fig. S16: Distribution of cancer-relevant perturbed edge categories across cancer types using NETMEDIUM.

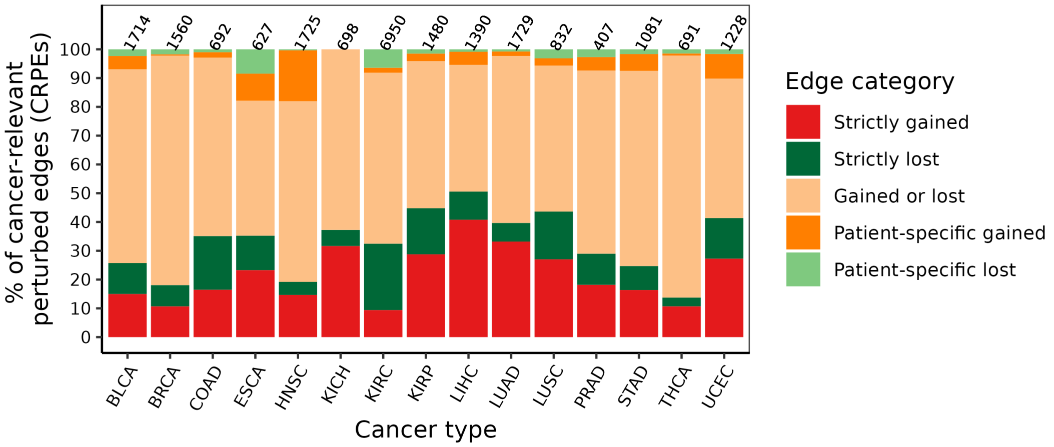


Supplementary Fig. S17: Distribution of cancer-relevant perturbed edge categories across cancer types using NETLOW.

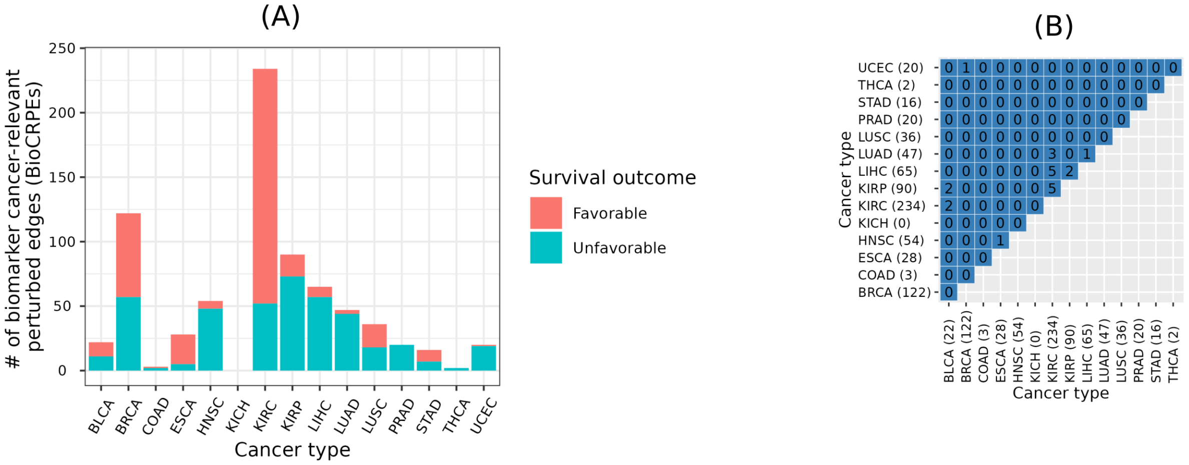


Supplementary Fig. S18: Biomarker cancer-relevant perturbed edges (CRPEs) across cancer types with respect to NETMEDIUM. (A) Number of BioCRPEs in each cancer type. (B) Counts of BioCRPEs shared between cancer types. Given two cancer types, $p$-values of the overlap between their BioCRPEs with respect to the background set of edges in a global exon-exon interaction network were calculated using the hypergeometric test and corrected using the Benjamini-Hochberg procedure to obtain the corresponding q-values. q-values less than or equal to 0.05 were considered significant. Numbers in each cell denote the counts of overlapping BioCRPEs between the corresponding cancer types.

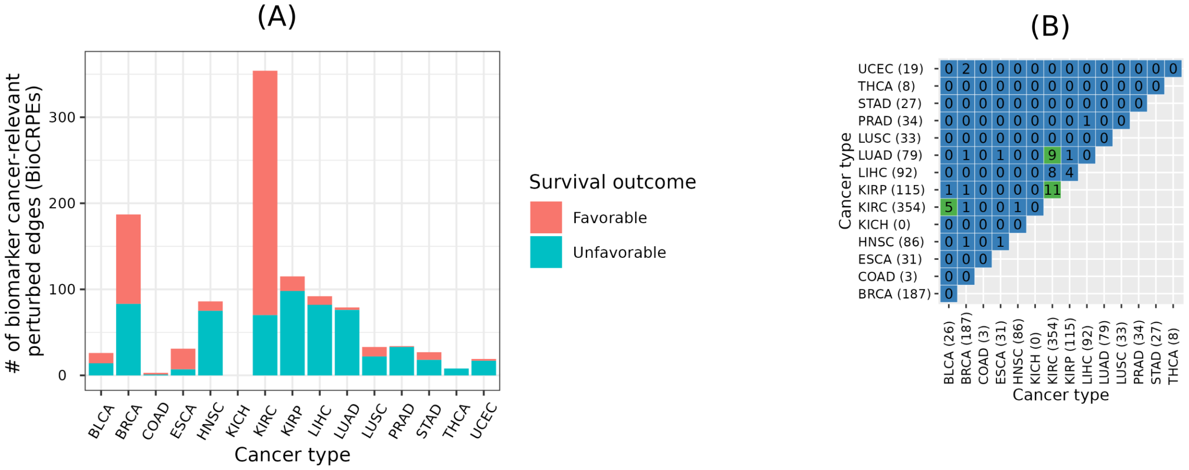


Supplementary Fig. S19: Biomarker cancer-relevant perturbed edges (CRPEs) across cancer types with respect to NETLOW. (A) Number of BioCRPEs in each cancer type. (B) Counts of BioCRPEs shared between cancer types. Given two cancer types, q-values of the overlap between their BioCRPEs were computed as for the overlaps between CRPEs in Supplementary Fig. S18. The green color cell indicates that the corresponding pair of cancer types show significantly high overlap between their BioCRPEs, while the blue color represents that the overlap was as expected by chance.

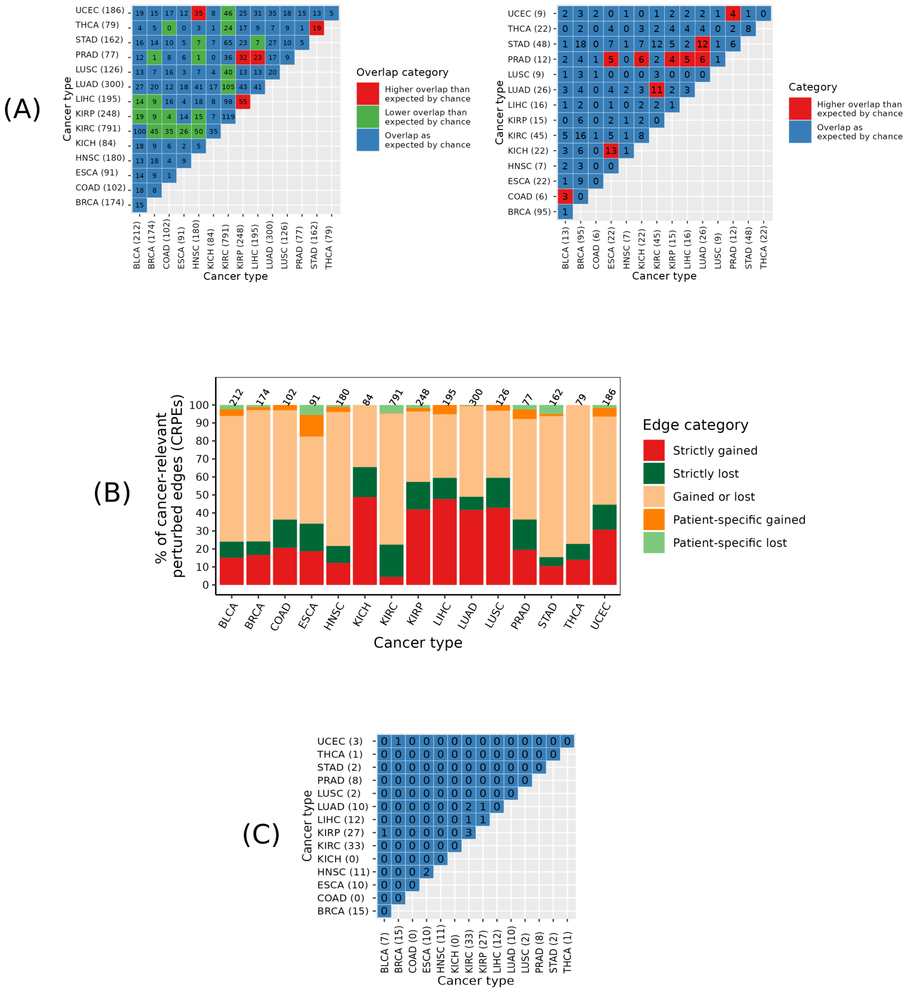


Supplementary Fig. S20: Key results corresponding to the Sections 3.4 to 3.7 of the main paper using an alternative definition of contact-based EEIs, where we defined a pair of exons to interact if and only if they have at least five residue-residue contacts as per our residue-residue definition in Section 2.3 of the main paper. All results are shown for NETHIGH created using the new contact-based EEI definition. (A) Left: Counts of CRPEs shared between cancer types, which can be interpreted similar to as Figure 7 of the main paper. Right: Numbers of enriched KEGG pathways shared between cancer types, which can be interpreted similar to as Figure 8 of the main paper. (B) Distribution of cancer-relevant perturbed edge categories across cancer types. (C) Pairwise overlaps between cancer types in terms of BioCRPEs, which can be interpreted similar to as Figure 10 of the main paper.

(A)

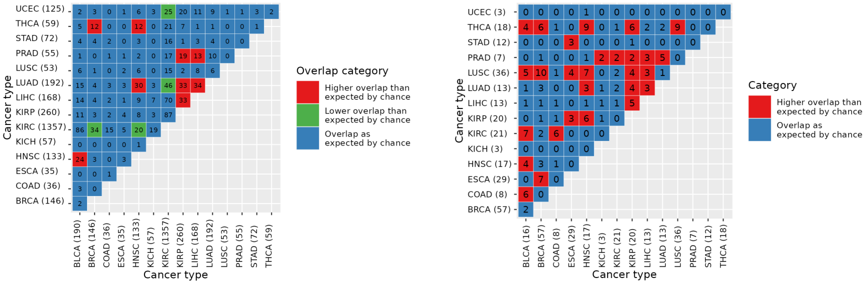


(B)

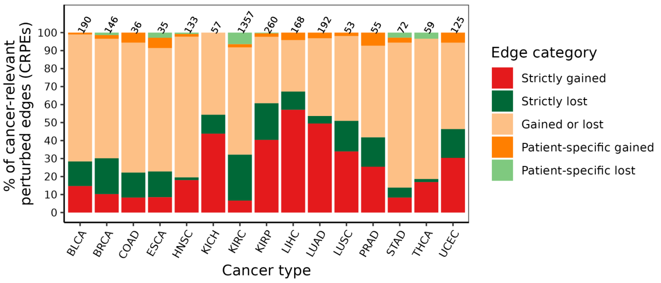


(C)

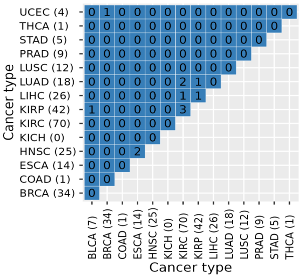


Supplementary Fig. S21: Key results corresponding to the Sections 3.4 to 3.7 of the main paper using an alternative survival analysis $p$-value cutoff of 0.01. All results are shown for NETHIGH. (A) Left: Counts of CRPEs shared between cancer types, which can be interpreted similar to as Figure 7 of the main paper. Right: Numbers of enriched KEGG pathways shared between cancer types, which can be interpreted similar to as Figure 8 of the main paper. (B) Distribution of cancer-relevant perturbed edge categories across cancer types. (C) Pairwise overlaps between cancer types in terms of BioCRPEs, which can be interpreted similar to as Figure 10 of the main paper.

### III Supplementary Tables

Supplementary Table S1: Cancer relevant perturbed edges (CRPEs) corresponding to (A) NETHIGH, (B) NETMEDIUM, and (C) NETLOW.

Supplementary Table S2: Cancer relevant perturbed edges (CRPEs) in partially perturbed protein complexes corresponding to (A) NETHIGH, (B) NETMEDIUM, and (C) NETLOW.

Supplementary Table S3: KEGG pathways significantly enriched in cancer relevant perturbed edges (CRPEs) corresponding to (A) NETHIGH, (B) NETMEDIUM, and (C) NETLOW.

Supplementary Table S4: Biomarker cancer relevant perturbed edges (BioCRPEs) corresponding to NETHIGH.

[2] Yangjun Zhang, Xiangyang Yao, Hui Zhou, Xiaoliang Wu, Jianbo Tian, Jin Zeng, Libin Yan, Chen Duan, Haoran Liu, Heng Li, Ke Chen, Zhiquan Hu, Zhangqun Ye, and Hua Xu. OncoSplicing: an updated database for clinically relevant alternative splicing in 33 human cancers. *Nucleic Acids Research*, 50(D1):D1340–D1347, 09 2021.
